# Engineering the substrate specificity of toluene degrading enzyme XylM using biosensor XylS and machine learning

**DOI:** 10.1101/2022.10.27.513980

**Authors:** Yuki Ogawa, Yutaka Saito, Hideki Yamaguchi, Yohei Katsuyama, Yasuo Ohnishi

**Affiliations:** Department of Biotechnology, Graduate School of Agricultural and Life Sciences, The University of Tokyo, Tokyo 113-8657, Japan; Collaborative Research Institute for Innovative Microbiology, The University of Tokyo, Tokyo 113-8657, Japan; Artificial Intelligence Research Center, National Institute of Advanced Industrial Science and Technology (AIST), Tokyo 135-0064, Japan; AIST-Waseda University Computational Bio Big-Data Open Innovation Laboratory (CBBD-OIL), Tokyo 169-8555, Japan; Department of Computational Biology and Medical Sciences, Graduate School of Frontier Sciences, The University of Tokyo, Chiba 277-8561, Japan

**Keywords:** XylM, enzyme engineering, directed evolution, machine learning, biosensor

## Abstract

Enzyme engineering using machine learning has been developed in recent years. However, to obtain a large amount of data on enzyme activities for training data, it is necessary to develop a high-throughput and accurate method for evaluating enzyme activities. Here, we examined whether a biosensor-based enzyme engineering method can be applied to machine learning. As a model experiment, we aimed to modify the substrate specificity of XylM, a rate-determining enzyme in a multistep oxidation reaction catalyzed by XylMABC in *Pseudomonas putida*. XylMABC naturally converts toluene and xylene to benzoic acid and toluic acid, respectively. We aimed to engineer XylM to improve its conversion efficiency to a non-native substrate, 2,6-xylenol. Wild-type XylMABC slightly converted 2,6-xylenol to 3-methylsalicylic acid, which is the ligand of the transcriptional regulator XylS in *P. putida*. By locating a fluorescent protein gene under the control of the *Pm* promoter to which XylS binds, a XylS-producing *Escherichia coli* strain showed higher fluorescence intensity in a 3-methylsalicylic acid concentration-dependent manner. We evaluated the 3-methylsalicylic acid productivity of XylM variants using the fluorescence intensity of the sensor strain as an indicator. The obtained data provided the training data for machine learning for the directed evolution of XylM. Two cycles of machine learning-assisted directed evolution resulted in the acquisition of XylM-D140E-V144K-F243L-N244S with 15 times higher productivity than wild-type XylM. These results demonstrate that an indirect enzyme activity evaluation method using biosensors is sufficiently quantitative and high-throughput to be used as training data for machine learning. The findings expand the versatility of machine learning in enzyme engineering.

## Introduction

With the recent development of machine learning, protein engineering research applying this technology has become increasingly widespread.^1^ In the directed evolution^2^ which is a method for protein engineering, the optimal sequence to obtain a certain protein function is searched for by screening. In this case, it is impossible to obtain comprehensive information regarding the relationships between the sequences and functions of all variants. Machine learning can learn the relationship between input data (sequence information of variants) and output data (functional information of protein variants) that is needed to construct a prediction model.^1^ In addition, the sequence space that can be explored by machine learning is much larger than the sequence space that can be covered by laboratory screening; therefore, the possibility of selecting the desired variants is higher.^1^ Thus, machine learning reduces the effort and cost of screening by efficiently narrowing the sequence space that needs to be explored. For example, Arnold *et al*. applied machine learning to the directed evolution of a putative nitric oxide dioxygenase from *Rhodothermus marinus*, which catalyzes carbon-silicon bond formation between phenyl-dimethyl silane and 2-diazopropanoate.^3^ Since this enzyme produces two possible enantiomers, the authors used machine learning-assisted directed evolution to engineer this enzyme to produce each of the two product enantiomers. Two rounds of evolution resulted in the identification of variants for selective catalysis with 93% and 79% enantiomeric excess.

A critical issue for the establishment of machine learning assisted directed evolution is the quantity of training data.^4^ Some methods, such as kernel-based methods, require dozens or hundreds of data points while other methods, such as deep neural networks, typically require thousands or even more data points. Thus, it is essential to develop a high-throughput enzyme evaluation system. Several studies have applied machine learning to directed evolution of proteins using various kinds of indicators for evaluating enzyme activities, such as fluorescence intensity,^5^ thermostability,^6^ cross-linking efficiency,^7^ or enantioselectivity.^8^ However, few studies have applied machine learning to improve protein function in practice. This is because the sequence and functional analysis of a large number of variants, which serve as training data, is labor-intensive.

Ligand-dependent transcriptional regulators are biological molecules that activate or repress the transcription of specific genes when they recognize small molecules inside or outside cells.^9^ By placing antibiotic resistance genes or fluorescent protein genes under the control of certain promoters to which ligand-dependent transcriptional regulators bind, such transcriptional regulators can be used as biosensors.^10^ Therefore, a transcriptional regulator that recognizes substrates or products of a target enzyme is useful for high-throughput and sensitive evaluation or selection of desired enzyme variant activities. Many researchers have used such screening systems to modify hosts or enzymes to improve the yield of desired products.^11–16^ In addition, the screening systems can be used to modify enzymes that are difficult to analyze *in vitro*, such as membrane proteins and those catalyzing multistep reactions.

Although a biosensor-based evaluation system is high-throughput, it is unclear whether the data acquired in this system would be sufficiently quantitative and accurate to be used as training data for machine learning. To the best of our knowledge, no study has explored the possibility of biosensor-based enzyme engineering using machine learning. In addition, it is unclear whether the output of a multistep enzyme cascade can accurately assess the activity of a target enzyme when the output is used as training data for machine learning. The multistep enzyme cascade provides some advantages for directed evolution. First, it can mitigate several issues, such as product inhibition and product toxicity.^4^ Second, it can avoid the requirement of chemically modified substrate analogs or bulky fluorescent groups, allowing enzyme variants to be screened on natural substrates. Third, it can offer opportunities to increase screening throughput by providing an output that can be evaluated at a higher speed and throughput than conventional chemical analyses, such as mass spectrometry and liquid chromatography.^4^ Thus, if it accurately reflects the activity of the target enzyme, the range of enzymes for which machine learning-assisted directed evolution is applicable would be extended.

XylS is a transcriptional activator of *Pseudomonas putida* that recognizes several aromatic compounds as ligands.^17^ XylS forms a complex with its ligand and binds to the *Pm* promoter as a dimer to induce transcription.^18,19^ We studied XylS for the development of biosensors for several aromatic compounds.^20,21^ In this study, we aimed to construct a new 2,6-xylenol degradation pathway using the biosensor XylS as a model experiment to show that biosensor-based and multi-step enzyme cascade-based screening methods would be applicable in machine learning-assisted directed evolution of enzymes. 2,6-Xylenol degradation pathways in *Mycobacterium* species,^22,23^ *Ochromonas danica*,^24^ and *Penicillium frequentans*^25^ have been reported. However, a method for engineering these pathways to improve degradation efficiency has not yet been developed. The development of a method for manipulating the degradation pathways of 2,6-xylenol-related xenobiotic compounds would provide a solution for constructing a new 2,6-xylenol degradation pathway. Here, we focused on the XylMABC system from *P. putida*, which involves a multi-step oxidation reaction that converts toluene and xylene into benzoic acid and *m-*toluic acid, respectively (Figure 1 (a)). After two cycles of machine learning-assisted directed evolution, we successfully acquired XylM variants with higher productivity than wild-type XylM. Our study demonstrates that machine learning-aided enzyme engineering using transcriptional regulators as biosensors should enable the rapid construction of novel aromatic compound degradation pathways.

**Figure 1.**
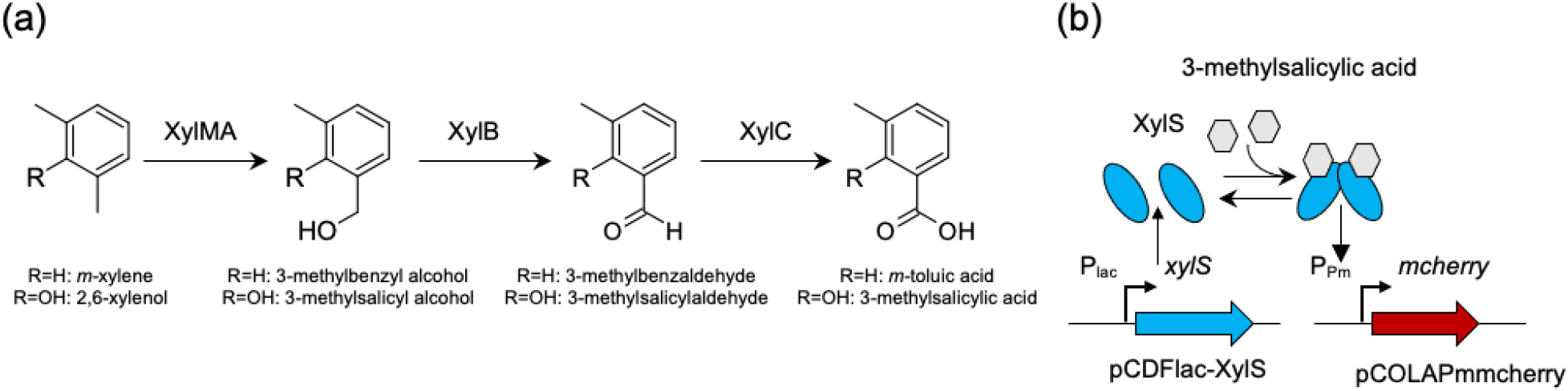
2,6-Xylenol detection system used in this study. (a) A multi-step oxidation reaction by XylMABC. (b) 3-Methylsalicylic acid detection by XylS using fluorescent protein gene.

## Results

### Construction of a screening system using XylS as biosensor

We evaluated the efficiency of the XylMABC system by detecting 3-methylsalicylic acid production from 2,6-xylenol using XylS as a biosensor. XylMABC are known to convert toluene and xylene to benzoic acid and *m-*toluic acid, respectively. We assumed that XylMABC also convert 2,6-xylenol into 3-methylsalicylic acid. In this reaction, XylMA catalyze the hydroxylation of one of the two methyl groups of 2,6-xylenol (XylM is the membrane-bound hydroxylase component; XylA is the reductase component)^26^, XylB oxidates 3-methylsalycilic alcohol to 3-methylsalicylaldehyde, and XylC oxidizes 3-methylsalicylaldehyde to 3-methylsalicylic acid, which is the ligand of XylS (Figure 1 (a)). By placing a fluorescence protein gene under the control of the *Pm* promoter, we constructed a system in which the host cell showed high fluorescence intensity in the presence of the XylS’s ligand (Figure 1 (b)). In this system, the fluorescence intensity of host cells was expected to increase in response to ligand concentration. We then examined the ligand detection range of XylS to confirm whether the fluorescence intensity of host cells was enhanced in a ligand concentration-dependent manner. pCDFlacXylS and pCOLAPmmcherry, which contain *xylS* under the control of the *lac* promoter and mCherry (a variant of red fluorescent protein) gene under the control of the *Pm* promoter, respectively^20^ (Figure 1 (b)), were introduced into *Escherichia coli* JM109. Recombinant cells were cultured in the presence of various concentrations of 3-methylsalicylic acid. A ligand concentration-dependent fluorescence response was clearly observed in the presence of 3-methylsalicylic acid at concentrations greater than 3 μM (Figure 2).

**Figure 2.**
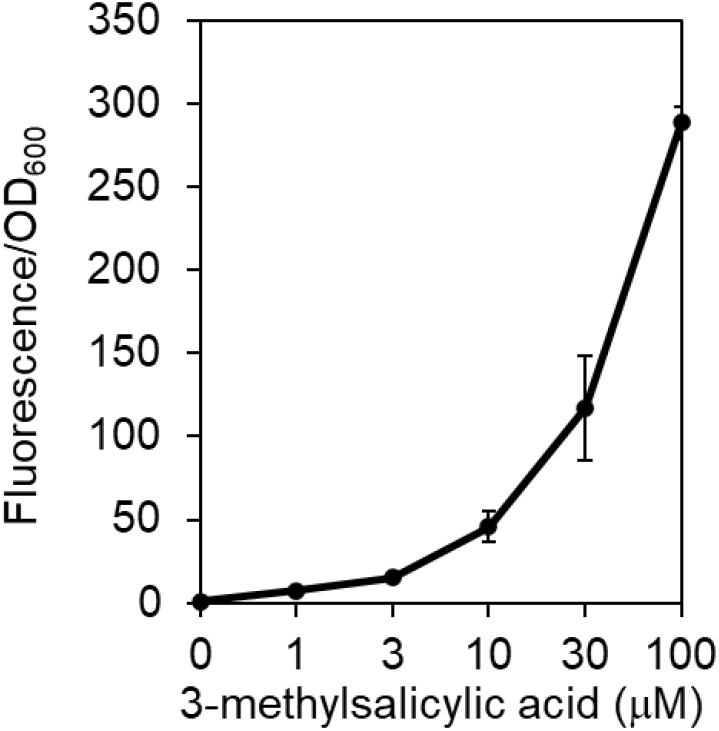
The fluorescence intensity of the XylS-producing sensor strain harboring mCherry gene under the control of *Pm* promoter in the presence of a variable concentration of 3-methylsalicylic acid. The error bars indicate standard error (n = 3).

### Substrate conversion efficiency of XylMABC

To analyze the substrate conversion efficiency of XylMABC, a bioconversion experiment was conducted using resting JM109-XylMABC cells. When a native substrate, *m-*xylene, was added, recombinant cells harboring XylMABC produced *m-*toluic acid with a high conversion rate (almost 80%) (Figure 3 (a)). In contrast, when the non-native substrate 2,6-xylenol was added, only a trace amount of 3-methylsalicylic acid was produced (conversion rate approximately 0.3%) (Figure 3 (b)). To determine the rate-limiting step, JM109-XylMABC cells were supplemented with the 3-methylsalicylic alcohol and 3-methylsalicylaldehyde intermediates. These compounds were converted to 3-methylsalicylic acid with high efficiencies (50% and 80%, respectively). The findings indicated that the initial reaction catalyzed by XylM was the rate-limiting step (Figure 3 (c)(d)). Therefore, we aimed to improve the substrate conversion efficiency of XylM to develop a new 2,6-xylenol degradation pathway.

**Figure 3.**
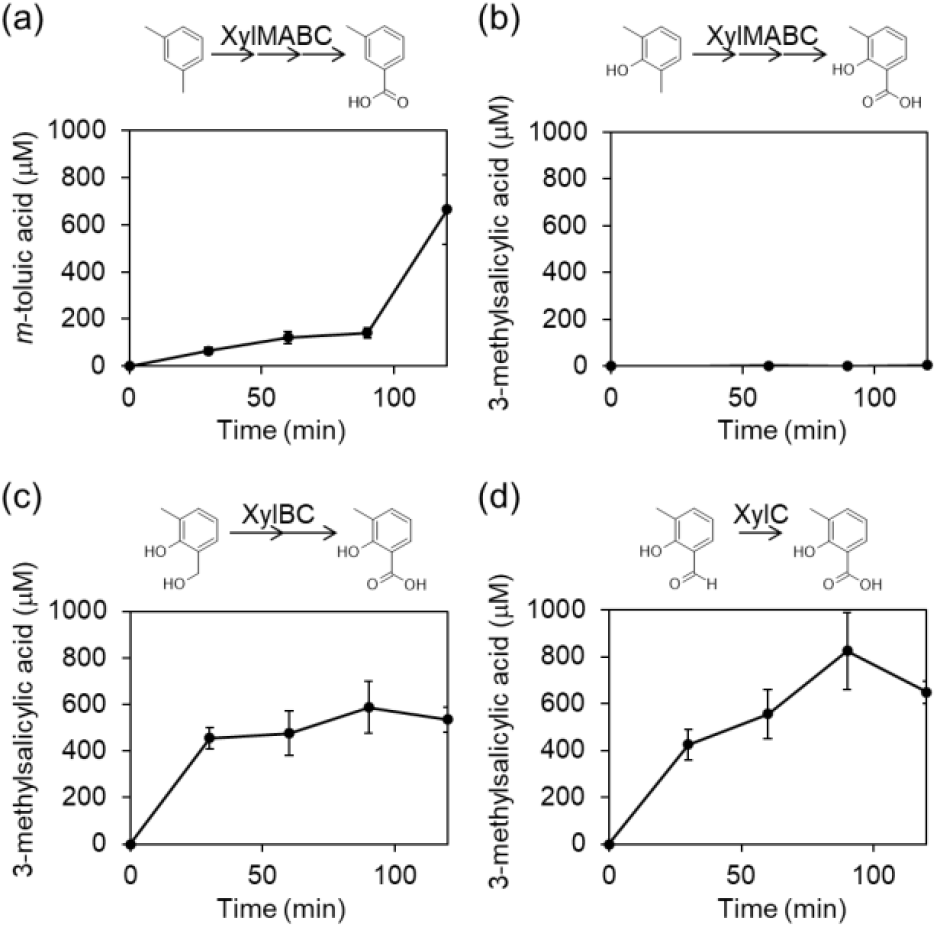
The substrate conversion efficiency of XylMABC analyzed by bioconversion using resting cells of JM109-XylMABC. (a) Conversion of *m-*xylene to *m-*toluic acid. (b) Conversion of 2,6-xylenol to 3-methylsalicylic acid. (c) Conversion of 3-methylsalicylic alcohol to 3-methylsalicylic acid. (d) Conversion of 3-methylsalicylaldehyde to 3-methylsalicylic acid. The error bars indicate standard error (n = 3).

### Selecting sites for mutagenesis based on modeled three-dimensional (3D) structure

The 3D structure of XylM was predicted using Phyre2^27^ based on the crystal structure of stearoyl-CoA desaturase (14% sequence identity with XylM).^28^ Alanine scanning of several residues located at the putative substrate-binding pocket was performed (Figure 4). Among these, T108A, H143A, D156A, and D158A completely abolished the activity of XylM, indicating that these amino acid residues are necessary for this activity (Figure 5). In contrast, D136A, P137A, D140A, V144A, L149A, Y167A, V240A, F243A, N244A, and I279A did not abolish the activity of XylM, although the production of *m*-toluic acid and/or 3-methylsalicylic acid was affected (the yields of these comounds increased or decreased to some extent) (Figure 5). Based on these results, five residues (P137, D140, V144, F243, and N244) that affected production and are located near the central iron coordination site of the putative substrate-binding pocket were selected as sites for mutagenesis.

**Figure 4.**
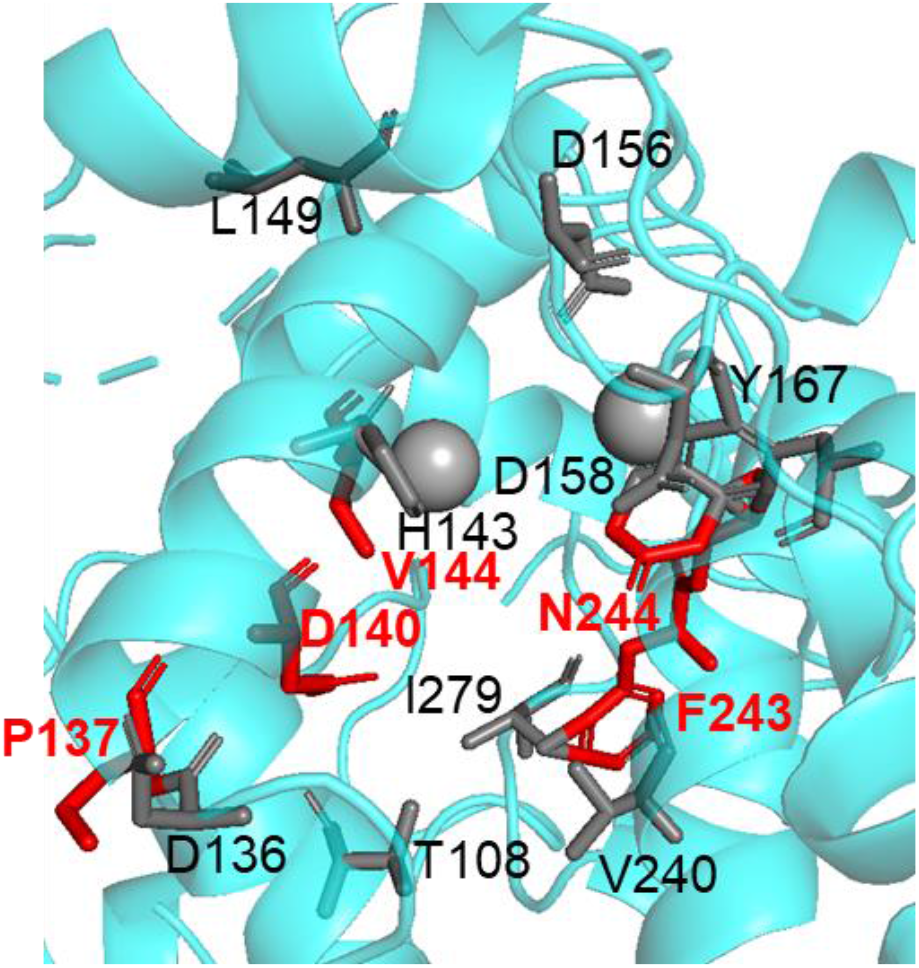
The putative substrate binding pocket of XylM modeled by Phyre2 based on the structure of a stearoyl-CoA desaturase (PDB ID: 4YMK). The amino acid residues surrounding the active site and selected for alanine scanning are shown in sticks. The amino acid residues which were selected for the mutagenesis are shown in red. Two spheres indicate the putative positions of ferrous ions.

**Figure 5.**
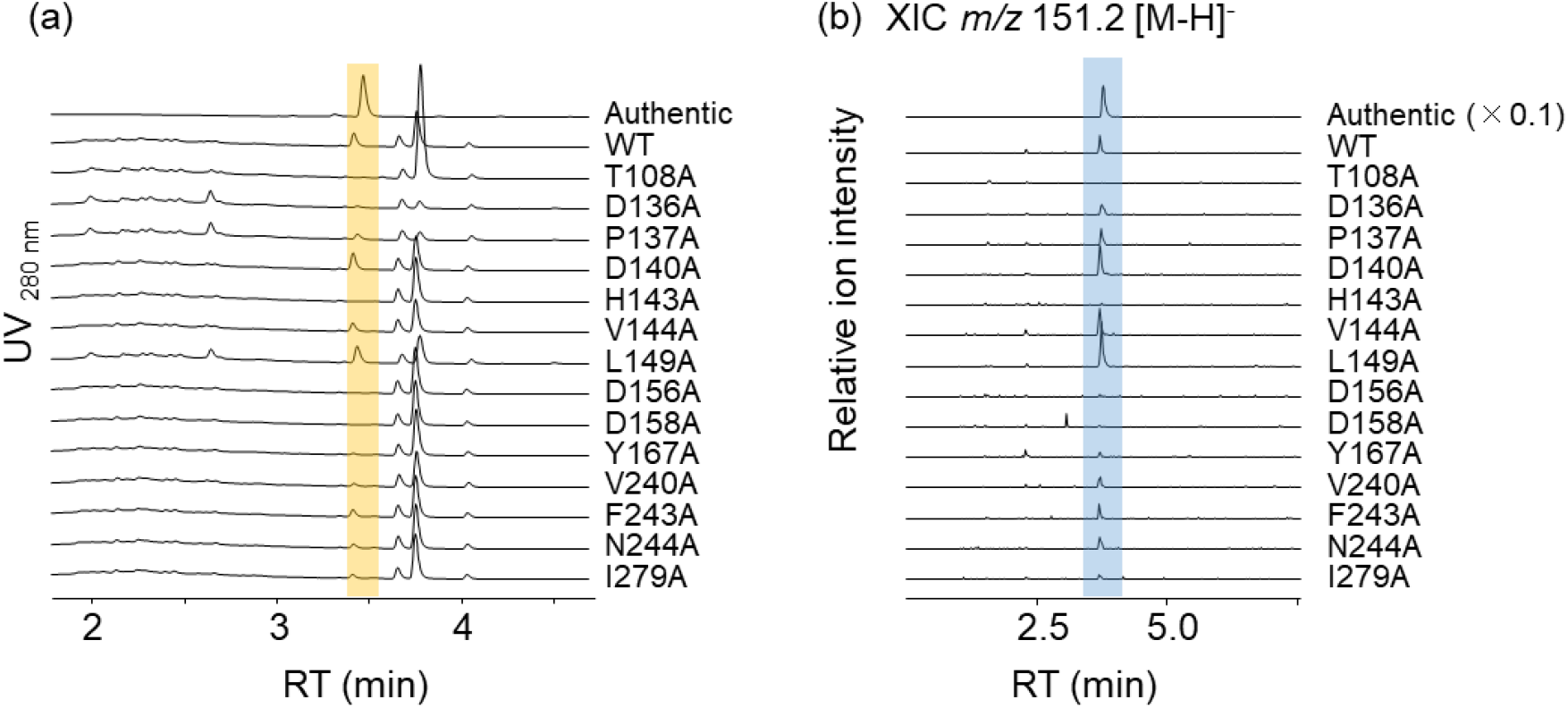
Alanine scanning of amino acid residues located at the putative substrate binding pocket of XylM. The activities of XylM variants were estimated by the resting cell bioconversion using JM109-XylMABC mutants. (a) The conversion of *m-*xylene to *m-*toluic acid. *m*-Toluic acid was detected by UV absorbance. “Authentic” indicates the chromatogram of *m-*toluic acid. (b) The conversion of 2,6-xylenol to 3-methylsalicylic acid. Because of the low yield, 3-methylsalicylic acid was detected by the liquid chromatography-mass spectrometry (LC-MS) analysis. Extracted ion chromatograms at [M–H]^−^ = *m/z* 151.2 are shown. “Authentic” indicates the chromatogram of 3-methylsalicylic acid.

### Directed evolution of XylM by machine learning

The XylM variant library obtained by simultaneous codon-randomization at five amino acid residues theoretically consists of 20^5^ = 3.2 × 10^6^ XylM variants. Because the number of XylM variants that can be evaluated by screening experiments is limited, we decided to screen the XylM variants using machine learning. In this method, we repeated the following processes twice: (i) training data preparation, (ii) prediction model training, (iii) activity prediction, and (iv) experimental evaluation of enzymatic activity. We constructed a screening system using a two-step culture method using two recombinant strains: a producer strain harboring XylMABC (JM109-pETlacXylMABC) and a sensor strain harboring plasmids containing *xylS* and the mCherry gene under the control of the Pm promoter (JM109-pCDFlacXylS-pCOLAPmmCherry). The sensor strain was inoculated into the culture supernatant of the producer strain after production, and incubated to estimate the yield of the product. We measured the fluorescence intensity of the sensor strain as the fluorescence intensity / optical density at 600 nm (OD_600_) of the cells, and defined it as the “fluorescence intensity activity.” This value was used to estimate the yield of 3-methylsalicylic acid. We trained the machine learning model to predict fluorescence intensity activity from the XylM variant sequence (described in Materials and Methods). The model is based on Bayesian optimization and considers various types of feature vectors.

### Preparation of the initial training data

For the acquisition of the initial training data, we constructed a library of 126 XylM variants that had substitution at five amino acid residues (P137, D140, V144, F243, and N244). This included all single amino acid-substituted XylM variants at five residues (19 amino acid substitution × five residues = 95 variants) and XylM variants with multiple amino acid substitutions at five residues. Hereafter, we describe the substitutions of P137, D140, V144, F243, and N244 as five letters. For instance, P137A, D140A, V144A, F243A, and N244S variants are described as AAAAS. The production of 3-methylsalicylic acid was evaluated by the ratio of the fluorescence intensity activity obtained from the strain with the XylM variant divided by that with wild-type XylM (WT-XylM) (Figure S1). We defined the resulting value as the relative fluorescence intensity activity of the XylM variant. Single substitutions of P137, D140, and F243 did not increase the relative fluorescence intensity activity. In contrast, the substitution of V144 resulted in several variants with improved relative fluorescence intensity activity (e.g., V144G). The N244S substitution drastically increased the relative fluorescence intensity activity. We also obtained six variants with multiple substitutions that showed an increased relative fluorescence intensity activity. Although these six variants had N244S substitutions, they had lower relative fluorescence intensity activity than XylM-N244S. This result indicates the difficulty of experimentally finding highly active variants with multiple substitutions. Therefore, we attempted to use machine learning with the obtained data to predict variants with higher activities.

### Machine learning and prediction of variants with better activities

Using the initial training data, we attempted to construct a machine learning model that predicts fluorescence intensity activity from the XylM variant sequence, as described in the Materials and Methods. As in our previous studies on machine learning-guided directed evolution^5,7^, we employed Bayesian optimization as a machine learning model. The prediction target was a fluorescence intensity score, defined as 1 – sigmoid (relative fluorescence intensity activity – 1). This score is high if the relative fluorescence intensity activity is higher than that of the wild-type enzyme, allowing the machine learning model to identify high-activity variants. For the feature vectors used in the machine learning model, we considered various amino acid descriptors based on physicochemical properties, chemical structures, evolutionary information, and their combinations. We performed benchmark experiments for feature selection and found that the combination of the ST-scale^29^ and position-specific score matrix (PSSM)^30^ achieved the best accuracy (denoted as ST-scale × PSSM, and detailed in the Materials and Methods and Figure S2 (a)(b)). In addition to these descriptor-based feature vectors, we also considered feature vectors based on representation learning by TAPE Transformer fine-tuned with XylM homologs^31,32^ (denoted as Evotuned-BERT). Evotuned-BERT achieved an accuracy comparable to ST-scale × PSSM in our benchmark experiment (see Materials and Methods and Figure S2 (c)). Therefore, we constructed two different machine learning models using either ST-scale × PSSM or Evotuned-BERT as feature vectors.

The constructed models were used to rank all XylM variants harboring multiple amino acid substitutions at the five selected residues (approximately 20^5^ XylM variants) using their fluorescence intensity scores predicted by machine learning. We extracted the top 100 variants predicted by the ST-scale × PSSM and Evotuned-BERT models independently and compared their sequence compositions (Figure S3). Intriguingly, the two models predicted similar but different sequences represented by “PXXFS” (Figure S3 (b)). Thirty-nine XylM variants were predicted by both models to be in the top 100 (Figure S4 (a)). In the ST-scale × PSSM model, two XylM variants were predicted to have higher scores than XylM-N244S, which showed the highest activity in the training data, while no XylM variants were predicted to have higher scores than XylM-N244S in the Evotuned-BERT model (Figure S4 (b)).

### Experimental evaluation of XylM variants predicted to have high activities

Of the top 100 XylM variants predicted by the ST-scale × PSSM and Evotuned-BERT models, we selected 50 variants for activity measurement. Twenty of these variants were selected from the above 39 variants commonly predicted by both models. Because most of them had sequences represented by PXXFS, their sequence diversity was low. To explore more diverse sequences, we selected the remaining 30 variants that were not represented by “PXXFS” and had substitutions at P137 or F243. The relative fluorescence intensity activity of the 50 selected XylM variants was evaluated (Figure S5). Remarkably, 19 XylM variants showed higher relative fluorescence intensity than XylM-N244S. Among them, the top five XylM variants with the highest relative fluorescence intensity activity (PEVLS, PNILS, PDLLS, PDILS, and PSDLS) were selected to evaluate their 3-methylsalicylic acid productivities. These XylM variants were expressed in recombinant *E. coli* along with XylABC, and 3-methylsalicylic acid production was measured when 2,6-xylenol was added to the growing cells. The cells all produced a higher amount of 3-methylsalicylic acid than did XylM-N244S (Figure 6). The PEVLS variant showed the highest productivity, with a 10-fold increase from WT-XylM and a 1.4-fold increase from XylM-N244S, the highest-activity variant in the training data.

**Figure 6.**
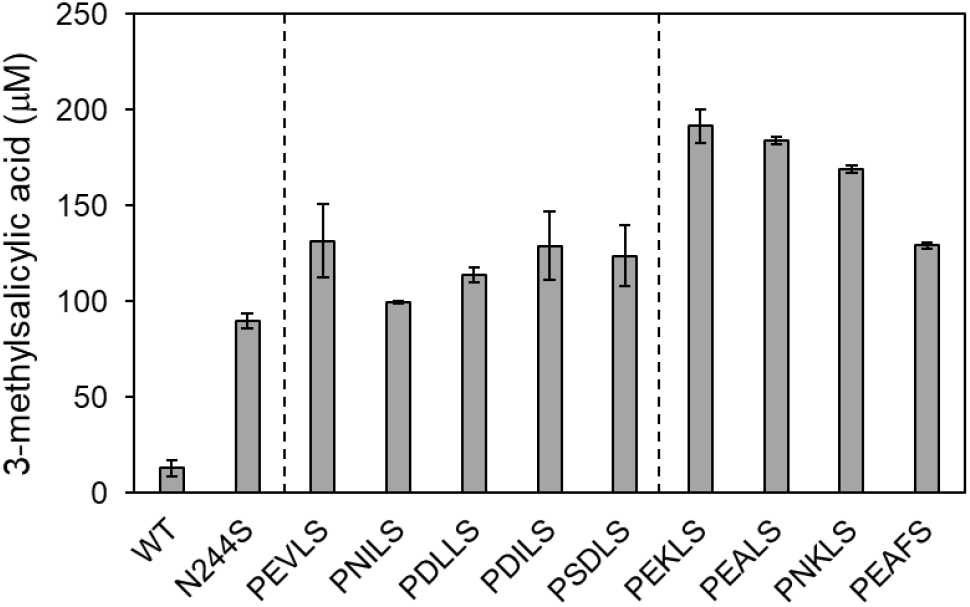
Evaluation of enzyme activity of the XylM variants predicted by machine learning. 3-Methylsalicylic acid production of the strains harboring the corresponding XylM variants and XylABC from 1 mM of 2,6-xylenol was determined during cultivation of cells. The error bars indicate standard error (n = 3).

### Second round of directed evolution using machine learning

The fluorescence intensity scores of the additional 50 XylM variants were added to the training data of machine learning to update both the models (ST-scale × PSSM and Evotuned-BERT models). The updated models were used to predict other XylM variants with high fluorescence intensity. The consensus sequence of the top 100 predicted XylM variants for both the ST-scale × PSSM and Evotuned-BERT models was P/C-XX-F/L-S (Figure S3 (c)). Forty-six XylM variants were commonly predicted by both models to be in the top 100 (Figure S6 (a)). Using the ST-scale × PSSM model, 85 XylM variants were predicted to have higher scores than XylM-PEVLS (the highest-activity variant in the first round) (Figure S6 (b)). On the other hand, 20 XylM variants were predicted to have higher scores than XylM-PEVLS when Evotuned-BERT was used. Of the top 100 XylM variants, we selected 50 for activity measurement. Similar to the first round, 19 of these XylM variants were selected for their high predicted scores by the ST-scale × PSSM and Evotuned-BERT models. Because XylM variants with P137C or P137S substitutions were predicted to have high scores, these XylM variants were additionally selected for activity measurement.

The relative fluorescence intensity activity of the 50 selected XylM variants was evaluated experimentally (Figure S7). Only one XylM variant, PEKLS, showed higher fluorescence intensity activity than XylM-PEVLS. The top four XylM variants with the highest relative fluorescence intensity activity (PEKLS, PEALS, PNKLS, and PEAFS) were selected to evaluate their 3-methylsalicylic acid productivities. These XylM variants were expressed in recombinant *E. coli* along with XylABC, and 3-methylsalicylic acid production was measured when 2,6-xylenol was added to the growing cells. The strains with XylM-PEALS and XylM-PNKLS produced 13- to 14-times higher concentrations of 3-methylsalicylic acid than WT-XylM (Figure 6). The strain with XylM-PEKLS produced 15-times higher amount of 3-methylsalicylic acid than WT-XylM.

### The substrate specificity of the acquired XylM variants

To analyze the substrate specificity of the acquired XylM variants, we measured the substrate conversion rates of WT-XylM, XylM-N244S, XylM-PEVLS, and XylM-PEKLS by resting cell bioconversion (Figure 7). We used resting cells for this experiment because the production of *m*-toluic acid from *m*-xylene could not be observed in the growing cells, probably due to the volatilization of *m*-xylene. When *m-*xylene was added as a substrate, the XylM-N244S, XylM-PEVLS, and XylM-PEKLS variants showed drastically reduced conversion rates compared to WT-XylM, although their conversion rates for 3-methylsalicylic acid were more than seven-times higher (Figure 7). These results indicate that the N244S substitution had a negative effect on the ability of XylM to recognize its native substrate, *m-*xylene, and a positive effect on its ability to recognize 2,6-xylenol. The other substitutions are likely to be important for further improvement of 2,6-xylenol recognition. Thus, we successfully engineered the substrate specificity of XylM.

**Figure 7.**
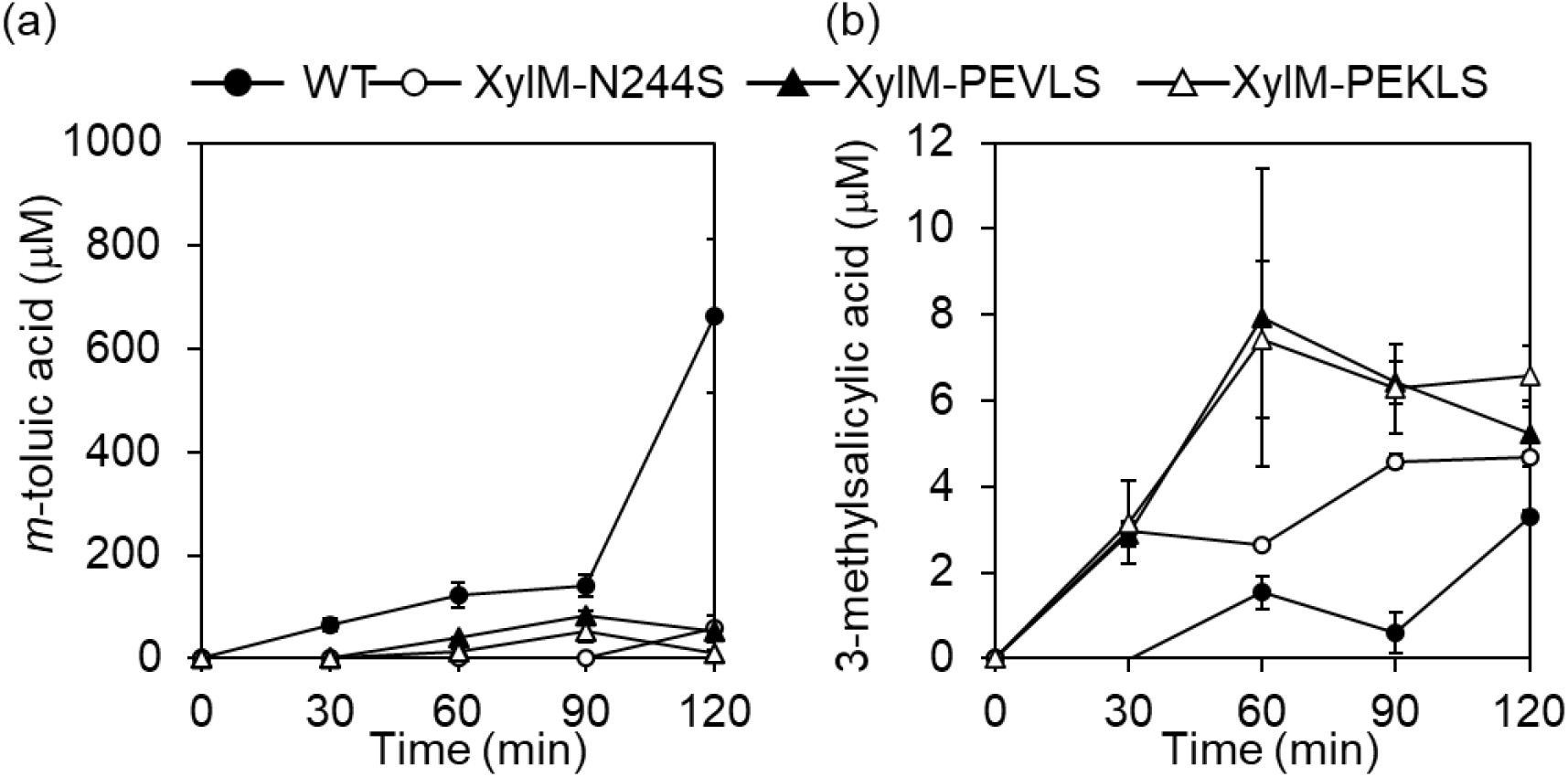
Evaluation of substrate specificity of the XylM variants predicted by machine learning. The substrate conversion by JM109 harboring the XylM variant and XylABC in the resting cell bioconversion were determined. (a) Conversion of *m-*xylene to *m-*toluic acid. (b) Conversion of 2,6-xylenol to 3-methylsalicylic acid. The error bars indicate standard error (n = 3).

## Discussion

Before the advent of machine learning-aided protein engineering, a typical conventional approach for designing variant libraries was to mimic amino acid substitutions observed in homologs. To investigate the advantages of machine learning, we surveyed amino acid substitutions in XylM homologs (Figure S8) and compared them with the machine learning-predicted XylM variants. Among the XylM homologs, P137 and D140 were highly conserved (Figure S8). Amino acid substitutions of these residues did not improve the 2,6-xylenol-oxidation activity in our experiments (Figure S1). In contrast, amino acid substitutions of V144, which is less conserved among XylM homologs (Figure S8), affected 2,6-xylenol-oxidation activity depending on the amino acid introduced (Figure S1). These results indicate that homolog information might provide some clues for designing single amino acid-substituted variants of XylM. However, this did not apply to multiple amino acid substitutions. For example, our first-round variant library designed using machine learning included XylM-CSLFS and XylM-CDIFS. Despite their amino acid substitutions at the conserved residue (P137C), they showed higher activity than XylM-N244S, suggesting a combinatorial effect of multiple amino acid substitutions. Furthermore, XylM-PEKLS, which showed the highest activity in our study, also had amino acid substitutions at the conserved residue (D140E). These results indicate that although single amino acid substitutions of P137 or D140 decrease 2,6-xylenol oxidation activity, combining them with other amino acid substitutions may increase the 2,6-xylenol oxidation activity. Thus, machine learning is effective for narrowing the sequence space of screening and for discovering unexpected highly active variants.

Another conventional approach is to introduce single amino acid substitutions near the active center based on structural information, accumulate substitutions that increase enzyme activity, and combine them to obtain a more active enzyme. However, this methodology is also expected to have difficulty obtaining highly active enzymes by combining multiple amino acid substitutions. In our analysis of single amino acid-substituted enzymes, only V144 and N244 substitutions were expected to contribute to increased 2,6-xylenol oxidation activity. However, to obtain the most active XylM variant (XylM-PEKLS) in our study, the D140E substitution seemed to be important, although the single D140E substitution reduced the activity. This observation further demonstrates that machine learning can be useful for improving enzyme activities.

As shown in Figure 7, N244S significantly reduced the activity of XylM toward its native substrate, *m-*xylene. This observation is consistent with the high conservation rate of N244 among XylM homologs (Figure S8), indicating its importance for recognizing *m-*xylene. The role of N244S in increasing 2,6-xylenol-oxidation activity was not clarified. However, we assumed that N244S substitution extended the space of the substrate-binding pocket to accommodate the hydroxy group of 2,6-xylenol, which is absent in *m*-xylene, and provides a hydrogen bond to recognize the additional hydroxy group.

The engineering of XylM to construct a new 2,6-xylenol degradation pathway serves as a model case for research on engineering the degradation pathway of aromatic compounds. Aromatic compounds are major sources of environmental pollution and it is important to detect and remove them from the environment. Many aromatic compound degradation pathways by microorganisms have been studied, but there are still many aromatic compounds that are difficult to efficiently degrade. Therefore, methods to modify known aromatic compound degradation pathways or construct new degradation pathways are required. However, few studies have addressed this issue.^33,34^ Because there are many naturally occurring transcriptional regulators that can detect various aromatic compounds, enzyme engineering using a product detection system based on transcriptional regulators combined with machine learning should enable the rapid construction of novel aromatic compound degradation pathways.

## Conclusion

In this study, we developed a novel enzyme engineering method by combining machine learning and a 3-methylsalicylic acid detection system using the biosensor XylS which was designed to control the expression level of a fluorescent protein. Several studies on enzyme engineering using machine learning have been reported^3,6,8^. However, whether biosensor-based indirect evaluation of enzymatic activity could be used as training data for machine learning has not been examined. This study shows that the final product of a multistep reaction pathway, including the target enzyme to be engineered, can be quantified indirectly *in vivo* using transcriptional regulator-dependent fluorescence protein expression. Such a quantification method is simple and has a high-throughput compared to methods using mass spectrometry or chromatography, and may be applicable to enzymes that are difficult to analyze *in vitro* because of their insolubility or physiological properties (e.g., membrane proteins such as XylM). This study also demonstrates that this biosensor-based indirect evaluation can be used as training data for machine learning. Significantly, these results expand the range of enzymes to which machine learning-aided engineering can be applied. Finally, this study provides a method to engineer aromatic compound degradation pathways by applying biosensors to detect certain aromatic compounds.

## Material and methods

### Strains, media, chemicals, and other materials

*E. coli* strain JM109 (Takara Biochemicals, Shiga, Japan) was used as the host for DNA manipulation and sensor strain construction. Luria-Bertani (LB) broth was purchased from Difco (Franklin Lakes, New Jersey, USA). All chemicals were purchased from FUJIFILM Wako Pure Chemical Corporation (Osaka, Japan), Sigma-Aldrich (St. Louis, Missouri, USA), and Tokyo Chemical Industry Co. Ltd. (Tokyo, Japan). The enzymes used for DNA manipulation were purchased from TaKaRa Bio Inc. (Shiga, Japan). Primers were purchased from Thermo Fisher Scientific (Waltham, Massachusetts, USA) and their sequences are listed in Table S1. The DNA sequences were analyzed by Fasmac Co. Ltd. (Kanagawa, Japan). *P. putida* mt-2 was purchased from National Institute of Technology and Evaluation (NITE).

### Construction of pETlacXylMAB-cat and pETlacXylMABC-cat plasmids

pETlacXylMAB-cat and pETlacXylMABC-cat contained *xylMAB* and *xylMABC*, respectively, under the control of *the lac* promoter. *P. putida* mt-2 (NBRC109109) was inoculated into 2 mL benzoic acid medium and pre-cultivated overnight at 30 °C. The overnight culture was then inoculated (1% v/v) into 2 mL of fresh benzoic acid medium and cultivated at 30 °C for an additional 48 h. The genome and pWW0 plasmid were extracted from the culture medium. A DNA fragment containing *xylMABC* was PCR-amplified with primers 1 and 2, using the extracted pWW0 as a template. The DNA fragment was digested with KpnI and PstI, and cloned into the corresponding sites of pCDFDuet-1, resulting in pCDFDuet-XylMABC. Unintentional mutations introduced during PCR were corrected by inverse PCR with primers 3 and 4, using pCDFDuet-XylMABC as a template, resulting in pCDFDuet-XylMABC-2. The template plasmid was then digested with DpnI. pCDFDuet-XylMABC-2 was digested with KpnI and PstI, and *xylMABC* was cloned into the corresponding sites of pETDuet-1, resulting in pETDuet-XylMABC. A DNA fragment containing the *lac* promoter was PCR-amplified with primers 5 and 6, using pCDFlac-1 as a template.^20^ The *lac* promoter was digested with NcoI and PstI, and cloned into the corresponding sites of pETDuet-XylMABC, resulting in pETlac-XylMABC. A DNA fragment containing the *cat* gene was PCR-amplified with primers 7 and 8 using pACYC184 as a template, and a DNA fragment containing the *xylMABC* genes under the control of the *lac* promoter was PCR-amplified with primers 9 and 10 using pETlacXylMABC as a template. Both amplified DNA fragments were connected using NEBuilder HiFi DNA Assembly Master Mix, resulting in pETlac-XylMABC-cat. A DNA fragment containing *xylMAB* under the control of the *lac* promoter was PCR-amplified with primers 11 and 12, using pETlac-XylMABC-cat as a template. The PCR product was self-cyclized during PCR and the template plasmid was digested with DpnI, resulting in pETlacXylMAB-cat.

### Measurement of the XylS detection range

*The E. coli* JM109 strain harboring pCDFlacXylS and pCOLAPmmcherry^20^ was pre-cultivated in LB medium at 37 °C overnight. Then, the overnight culture was inoculated (1% v/v) into 200 μL of LB medium containing 50 μM isopropyl-β-D-thiogalactopyranoside (IPTG), 50 mg/L kanamycin, 50 mg/L streptomycin, and 1, 3, 10, 30, or 100 μM 3-methylsalicylic acid and cultivated at 37 °C for 10 h. Cells were harvested by centrifugation for 15 min at 1100 × g and resuspended in 200 μL of PBS buffer (pH 7.4–7.6). The OD_600_ (optical density at 600 nm) and fluorescence (excitation wavelength of 587 nm and emission wavelength of 610 nm) values were measured in triplicate using a microplate reader Spectramax M2 (Spectramax M2, Molecular Devices, San Jose, California).

### Validation of XylMABC activity by resting cell bioconversion

*E. coli* JM109 strain harboring either pETlacXyl-MABC-cat or pETlacXylM*ABC-cat was pre-cultivated overnight at 37 °C. The overnight culture was inoculated (1% v/v) into 100 mL LB medium containing 34 mg/L chloramphenicol. The cells were cultivated at 37 °C for 2 h, and 1 mM IPTG was added to the culture. The cells were cultivated for another 4 h and harvested by centrifugation at 1100 × g for 15 min. The cells were resuspended in 14 mL potassium phosphate buffer and incubated at 30 °C with shaking at 200 rpm for 5 min. Then, the substrates (1 mM of *m-*xylene, 2,6-xylenol, 3-methylsalicyl alcohol, or 3-methylsalicylaldehyde) were added, and the cells were incubated for 2 h. A portion (1 mL) of culture medium was collected every 30 min, and the cells were harvested by centrifugation for 15 min at 1100 × g. The supernatant was filtered, and 5 μL of the supernatant was analyzed by LC-ESIMS analysis using a Shimadzu (Kyoto, Japan) Nexera-i LC-2040C 3D Plus equipped with a COSMOCORE 2.6 C18 packed column (2.1×100 mm, Nacalai Tesque, Kyoto, Japan) for LC-MS analysis. A Shimadzu LCMS-8040 instrument was used for MS analysis. A linear gradient of water-acetonitrile containing 0.1% formic acid at a flow rate of 0.6 mL/min (5-100% acetonitrile) was used as the solvent.

### Synthesis of 3-methylsalicylic alcohol

3-Methylsalicylaldehyde (1 mM) was dissolved in 10 mL methanol and kept on ice. NaBH_4_ (2 mM) was slowly added, and the mixture was stirred for 30 min. Subsequently, 1 M HCl (10 ml) was added to stop the reaction. The aqueous layer was extracted with ethyl acetate, combined with the organic layer, and washed with a saturated NaHCO_3_ solution and brine. The organic layer was dried over MgSO_4,_ filtered, and concentrated to obtain 3-methylsalicylic alcohol (yield: quant.). The obtained compound was confirmed by comparing the ^1^H NMR data with previously reported values. ^1^H NMR (chloroform, 500 MHz) 7.41 (s, 1H), 7.08 (d, J = 7.5, 1H), 6.86 (d, J = 6.5, 1H), 6.75 (t, 1H), 5.10 (s, 1H), 4.83 (s, 2H), 2.25 (s, 3H).

### Measurement of fluorescence intensity activities of XylM variants in growing cells

*E. coli* JM109 strain harboring each pETlacXylMABC-cat derivative was precultivated at 37 °C for 8 h, and the pre-culture was inoculated (1% v/v) into 200 μL of LB medium containing 3 mM 2,6-xylenol, 50 μM IPTG, and 34 mg/L chloramphenicol. The cells were cultivated for 16 h and harvested by centrifugation at 1100 × g for 10 min. Next, 120 μL of the supernatant was transferred to another microwell. Pre-cultivated *E. coli* strains harboring pCDFlacXylS and pCOLAPmmcherry were inoculated (1% v/v) into the supernatant. Next, 100 μM IPTG, 50 mg/L streptomycin, and 50 mg/L kanamycin at the final concentration were added to the supernatant. The cells were cultivated for 10 h and harvested by centrifugation at 1100 × g for 10 min. The cells were resuspended in 200 μL of PBS buffer (pH 7.4-7.6). The OD_600_ and fluorescence (excitation wavelength of 587 nm and emission wavelength of 610 nm) values were measured in triplicate using a microplate reader Spectramax M2 (Molecular Devices).

### Measurement of 3-methylsalicylic acid productivities of strains harboring XylM variants and XylABC in growing cells

*The E. coli* JM109 strain harboring each pETlacXylMABC-cat derivative was pre-cultivated at 37 °C overnight, and the pre-culture was inoculated (1% v/v) into 100 mL of LB medium containing 1 mM 2,6-xylenol, 50 μM IPTG, and 34 mg/L chloramphenicol. The cells were cultivated for 8 h, and 30 μL of the supernatant was centrifuged at 15000 rpm (20630 × g) for 5 min, filtered, and subjected to LC-ESI-MS analysis.

### Construction of single amino acid-substituted XylM variants

A single amino acid-substituted *xylM* library harboring mutations at the codon for a certain amino acid was constructed by inverse PCR using a pair of primers for amino acid substitution at the codon to be changed (primers 15-48, 52-92). The PCR product was self-cyclized, and the template plasmid was digested with DpnI.

### Construction of the five-residue-substituted XylM variants

Two DNA fragments were PCR-amplified with primers 94 and 30 and primers 49 and 44 using pETlacXylMABC-cat as a template to perform site-saturation mutagenesis at five amino acid residues, P137, D140, V144, F243, and N244. The DNA fragments were connected using NEBuilder HiFi DNA Assembly Master Mix. The resulting plasmids were used to transform JM109, resulting in a P137-D140-V144-F243-N244 mutated XylM library. pETlacXylM-F243L-N244S-XylABC-cat, pETlac-XylM-F243Y-N244S-XylABC-cat, and pETlacXylM-F243M-N244S-XylABC-cat were constructed by inverse PCR using primers 95 and 30, primers 96 and 30, and primers 97 and 30, respectively, using pETlacXylMABC-cat as a template. The PCR products were self-cyclized, and the template plasmid was digested with DpnI. Meanwhile, the plasmids encoding the triple or quadruple amino acid substituted XylM variants (e.g., pETlacXylM-P137A-D140A-F243Y/L/M-N244S), a DNA fragment which was PCR-amplified with a pair of primers (126 + 44, 98 + 44, and 98 + 16) using any of pETlacXylM*ABC-cat vectors (pETlacXylM-F243L-N244S-XylABC-cat, pETlac-XylM-F243Y-N244S-XylABC-cat, pETlacXylM-F243M-N244S-XylABC-cat, or pETlacXylM-N244S-XylABC-cat) as a template was connected with a primer (primers 105-125) acting as a single strand oligo DNA with single or double amino acid substitutions by NEBuilder Hifi DNA Assembly Master Mix. The template plasmid was then digested with DpnI. The plasmids encoding the double or triple amino acid-substituted XylM variants (e.g., pETlacXylM-V144G-N244S-XylABC-cat and pETlacXylM-P137C-V144G-N244S-XylABC-cat) were constructed by inverse PCR with a pair of primers (32 + 51, 95 + 30, 96 + 27, 97 + 30, 96 +30, 95 + 30, and 32 + 51) using any of pETlacXylM*ABC-cat vectors encoding single amino acid-substituted XylM variants as a template.

To construct a series of pETlacXylM*ABC-cat vectors used in the experiments after the first machine learning, inverse PCR was performed using appropriate sets of primers 101-107, 109-110, 115, and 128-147 (template: pETlacXylM-F243L-N244S-ABC-cat) and 128, 133-137, 141, 145, and 148-153 (template: pETlacXylM-N244S-ABC-cat). After inverse PCR, each PCR product was self-cyclized, and the template plasmid was digested with DpnI.

All of the constructed *xylM* sequences were determined using primers 13 and 14 to confirm correct mutations.

### Machine learning model

As in our previous studies on the machine learning-guided directed evolution of proteins,^5,7^ Bayesian optimization was employed as a machine learning model. We used the Bayesian optimization software COMBO^35^, which implements a Gaussian process regressor based on a linear model using a random feature map as follows:

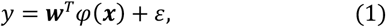

where *y* is the fluorescence intensity score of the protein (defined in the next section), ***x*** is the feature vector of the protein, *φ*(***x***) is a random feature map (with a dimensionality of 5,000), and *ε* is an error term. A radial basis function kernel was used as a kernel function with hyperparameters optimized using the type-2 maximum likelihood method implemented in COMBO. Given a training dataset {(*y*, ***x***)}, COMBO fits a weight vector ***w*** such that the fluorescence intensity score ***y*** can be predicted from the feature vector ***x***. For a given variant, COMBO can compute the acquisition function by Thompson sampling,^35^ which evaluates the probability that the fluorescence intensity score of the variant is higher than that of any variant in the training dataset. These values were used to rank all the possible variants.

In this study, we performed two rounds of machine learning. In the first round, the initial training data were prepared (Figure S1), and two separate machine learning models were constructed based on different feature vectors (ST-scale ×PSSM and Evotuned-BERT, described in the next section). These models were used to rank all possible variants and to obtain two ranking lists (Data S1). In the second round, the top-ranked variants from first-round machine learning were used as additional training data to update the models and ranking lists (Data S2).

### Fluorescence intensity score

We set the aim of machine learning to discover improved variants with higher relative fluorescence intensity activity (a measure of enzyme activity). For this purpose, the following fluorescence intensity score was used in the machine learning model:

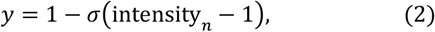

where *σ* (·) is a sigmoidal function and intensity_*n*_ is the relative fluorescence intensity activity. This score takes a high value if the relative fluorescence intensity activity of a variant is higher than that of the wild-type enzyme, allowing the machine learning model to identify high-activity variants.

### Feature vectors based on amino acid descriptors

The feature vectors considered in this study are broadly divided into two categories: amino acid descriptors and representation learning. Amino acid descriptors are feature vectors that are defined for each type of amino acid. We constituted a feature vector ***x*** of the XylM protein by concatenating the feature vectors of the amino acids at the five mutated residues. For a feature vector of each amino acid, we considered a variety of amino acid descriptors including ST-scale^29^, Z-scale,^36^ T-scale,^37^ FASGAI,^38^ MS-WHIM,^39^ ProtFP,^40^ VHSE,^41^ and BLOSUM-based features.^42^ For the characteristics of these descriptors, see the review by van Westen et al.^40^ In addition, PSSM was used as a feature vector to incorporate the evolutionary information from XylM homologs. Specifically, we constructed PSSM by a PSI-BLAST^30^ iterative homology search using the wild-type XylM sequence as a query against the nr database. The homology search was iterated five times, with 200 homologs used to update the PSSM per iteration. This produced a PSSM {*s*_*ij*_} for each residue *i* and amino acid *j*. The PSSM-based feature vector of the XylM protein was constructed by concatenating the PSSM values of the five mutated residues and their amino acids.

To select the optimal feature vector, a feature selection procedure was performed, as in our previous studies.^5,7^ Briefly, we performed a benchmark experiment in which COMBO was set to find XylM-N244S (the highest-activity variant) from the initial training data using a Bayesian optimization procedure (Figure S2 (a)). In this experiment, the ST-scale achieved better results than the other descriptors in terms of the amount of training data needed to find XylM-N244S. The ST-scale was further combined with the PSSM by concatenating the feature vectors, achieving slightly better results (Figure S2 (b)). Therefore, we used the combination of the ST-scale and PSSM as the feature vector for our descriptor-based model in all other parts of this study (denoted as ST-scale ×PSSM).

### Feature vectors based on representation learning

In addition to the amino acid descriptors, we considered feature vectors based on representation learning. Representation learning is a method for learning the feature representation of a protein using a large number of unlabeled amino acid sequences. We used the representation learning model TAPE Transformer^31^, which was pre-trained on the entire Pfam database. To incorporate the evolutionary information of XylM, we fine-tuned the pre-trained TAPE Transformer using XylM homologs via a method proposed previously^32^ (denoted as Evotuned-BERT). The performance of Evotuned-BERT was evaluated using the same benchmark experiment as that of amino acid descriptors, and the results were comparable to those of ST-scale×PSSM (Figure S2 (c)). Thus, we decided to use the machine learning model based on Evotuned-BERT separately from that based on the amino acid descriptors.

## Supporting Information

The relative fluorescence intensity activities of the XylM variant library used for training data for the first machine learning; benchmark of feature vectors; sequence composition of the XylM variants predicted by machine learning; first-round top 100 prediction using ST-scale×PSSM score or Evotuned-BERT score; the relative fluorescence intensity activities of XylM variants of first-round library; second-round top 100 prediction using ST-scale × PSSM score or Evotuned-BERT score; the relative fluorescence intensity activities of XylM variants of second-round library; multiple sequence alignment of 96 XylM homologs at P137, D140, V144, F243, and N244; primer list used in this study (PDF)

Ranking list by ML prediction using the initial XylM variant library as training data; ranking list by ML prediction using the second-round XylM variant library as additional training data. This list was used to design the third-round XylM variant library (xlsx)

## AUTHOR INFORMATION

### Author Contributions

Y.K., Y.Oh., and Y.S. designed the study. Y.Og. performed biochemical experiments. Y.Og. and Y.S. analyzed the data. Y.S. and Y.H. performed the machine learning. Y. Og., Y. K., Y. S., and Y. Oh. wrote the paper. All authors have read and approved the final manuscript. Y.K., Y.S., and Y.Oh. supervised the study. Y. Og. and Y. S. contributed equally.

## Notes

The authors declare no competing financial interests.

## ACKNOWLEDGMENTS

This research was supported in part by Grants-in-Aid for Challenging Exploratory Research (16K14880 to Y.K.) and JSPS Fellows (20J12216 to Y.Og.) from the Japan Society for the Promotion of Science (JSPS), the JACI Prize for Encouraging Young Researcher (to Y.S.), and the JSPS A3 Foresight Program (to Y. Oh.). Computations were partly performed on the NIG supercomputer at ROIS National Institute of Genetics, and AI Bridging Cloud Infrastructure (ABCI) at National Institute of Advanced Industrial Science and Technology (AIST). We thank Dr. Hikaru Nakazawa and Dr. Mitsuo Umetsu at Tohoku University for their useful advice regarding this study.

