## Supplementary Information for "Engineering the substrate specificity of toluene degrading enzyme XylM using biosensor XylS and machine learning"

AUTHOR ADDRESS

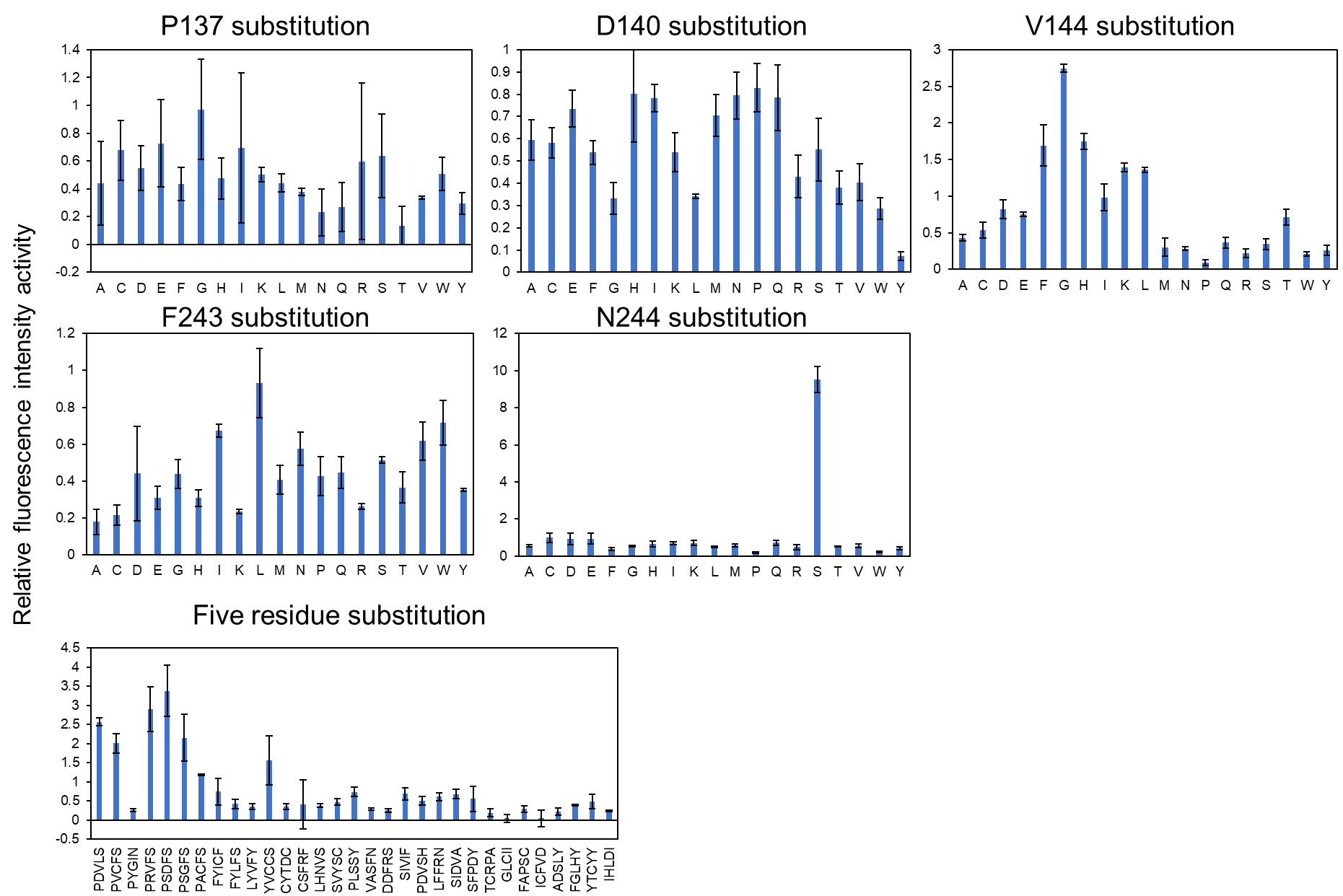

**Figure S1.** The relative fluorescence intensity activities of XylM variants library used for training data for the first machine learning. The error bars indicate standard error (n = 3).

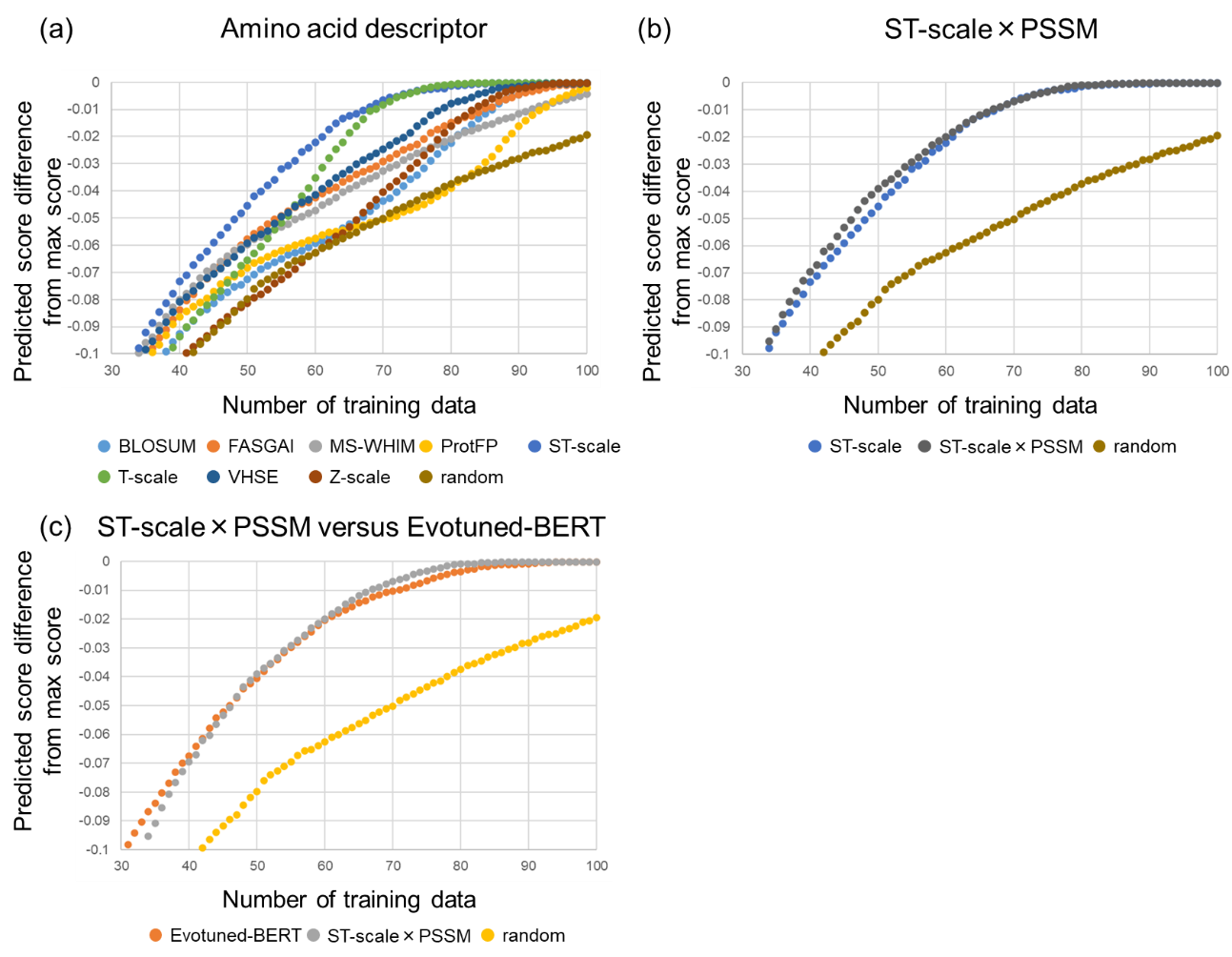

**Figure S2.** Benchmark of feature vectors. (a) For each type of feature vector, Bayesian optimization was conducted to find variants with high fluorescence intensity scores among the initial XylM variant library. The fluorescence intensity score of the best variant found is plotted against an increasing number of training data. The Bayesian optimization procedures were repeated 1,000 times with different choices of initial training data points (N=5), and the average results are shown. Feature vectors achieving higher fluorescence intensity scores at smaller numbers of training data are regarded as good feature vectors. ST-scale showed better performance than the other feature vectors. (b) ST-scale×PSSM showed slightly better performance than ST-scale only. (c) Evotuned-BERT and ST-scale×PSSM showed the comparable performance with each other.

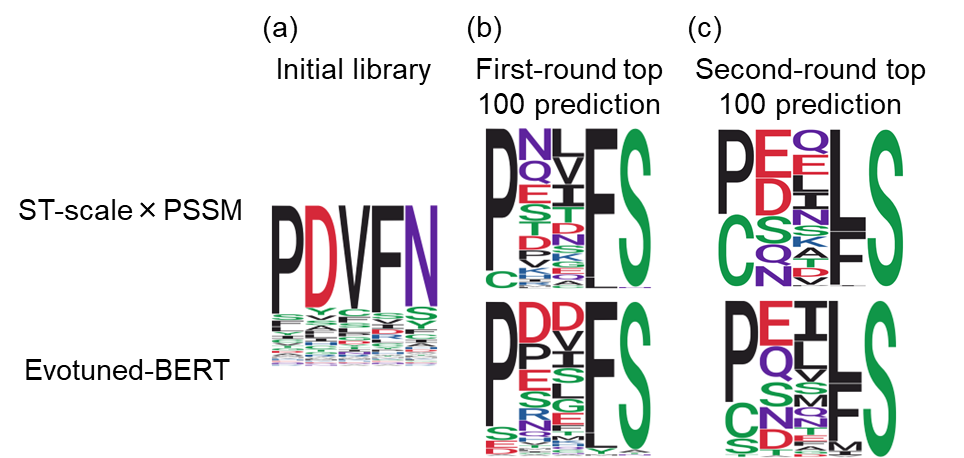

**Figure S3.** Sequence compositions of the XylM variants predicted by machine learning. In each case, the top 100 predicted variants are shown. The wild-type sequence is PDVFN. Sequence logo representation of amino acids at the five mutated residues of (a) initial training data, (b) first-round top 100 prediction, and (c) second-round top 100 prediction.

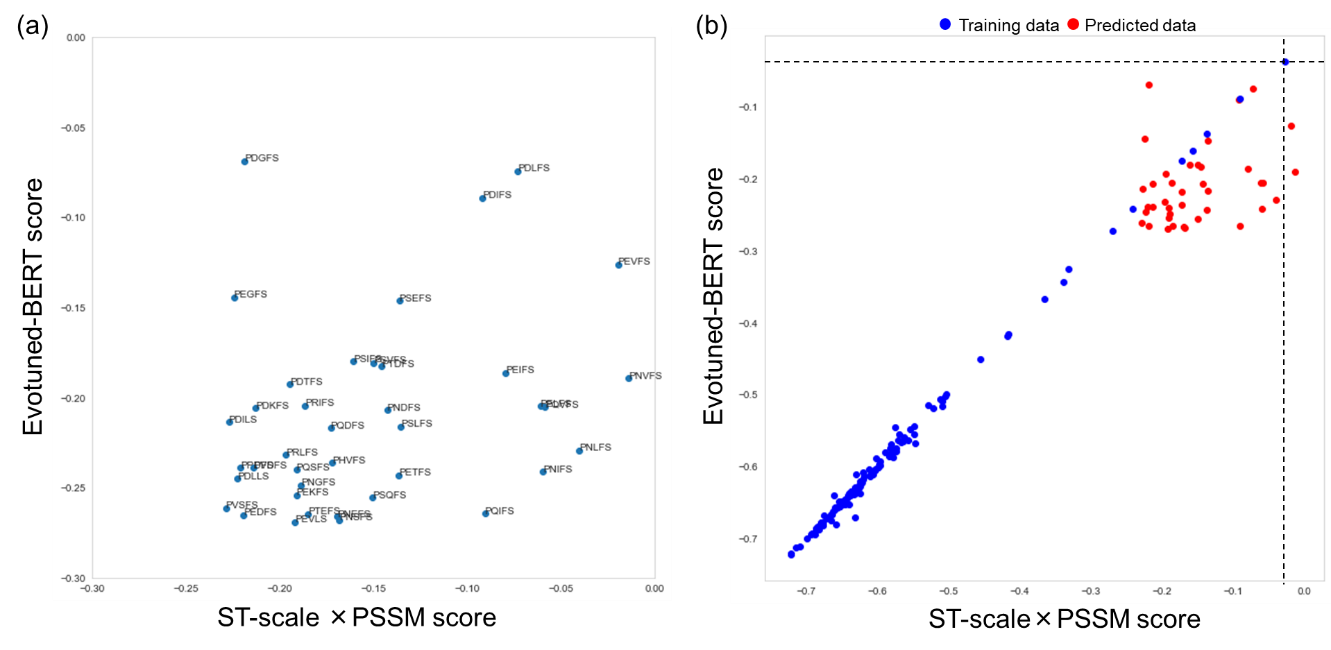

**Figure S4.** Consensus analysis of the first-round top 100 variants predicted by ST-scale×PSSM and Evotuned-BERT models. Predicted scores by the two models are shown. (a) 39 XylM variants commonly predicted by both models to be in the top 100. (b) Comparison of XylM variants of the initial library (training data) and the first-round top 100 prediction. The highest-activity variant in the training data (XylM-N244S) is shown with dashed lines.

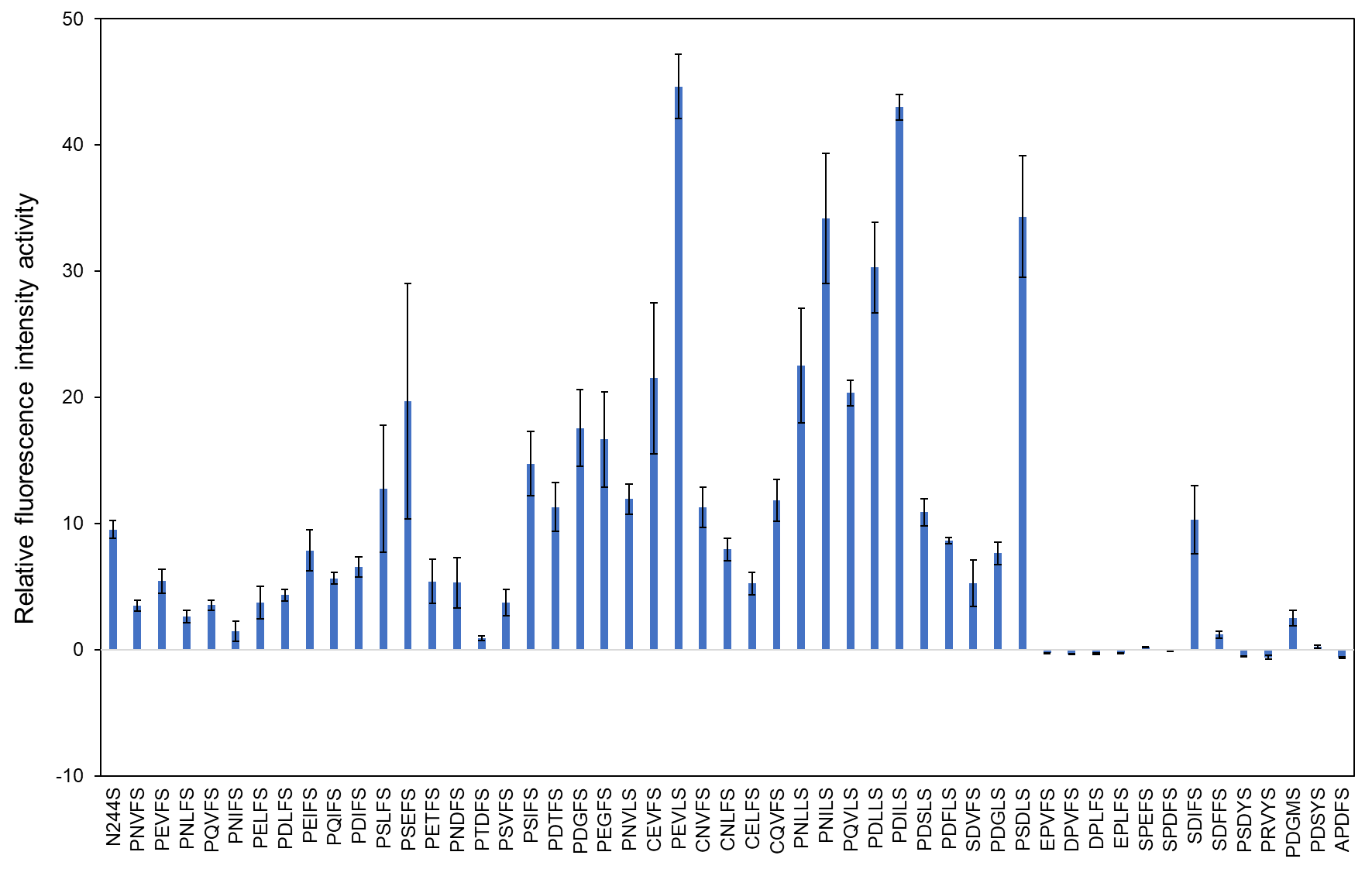

**Figure S5.** The relative fluorescence intensity activities of XylM variants of the first-round library. The error bars indicate standard error (n = 3).

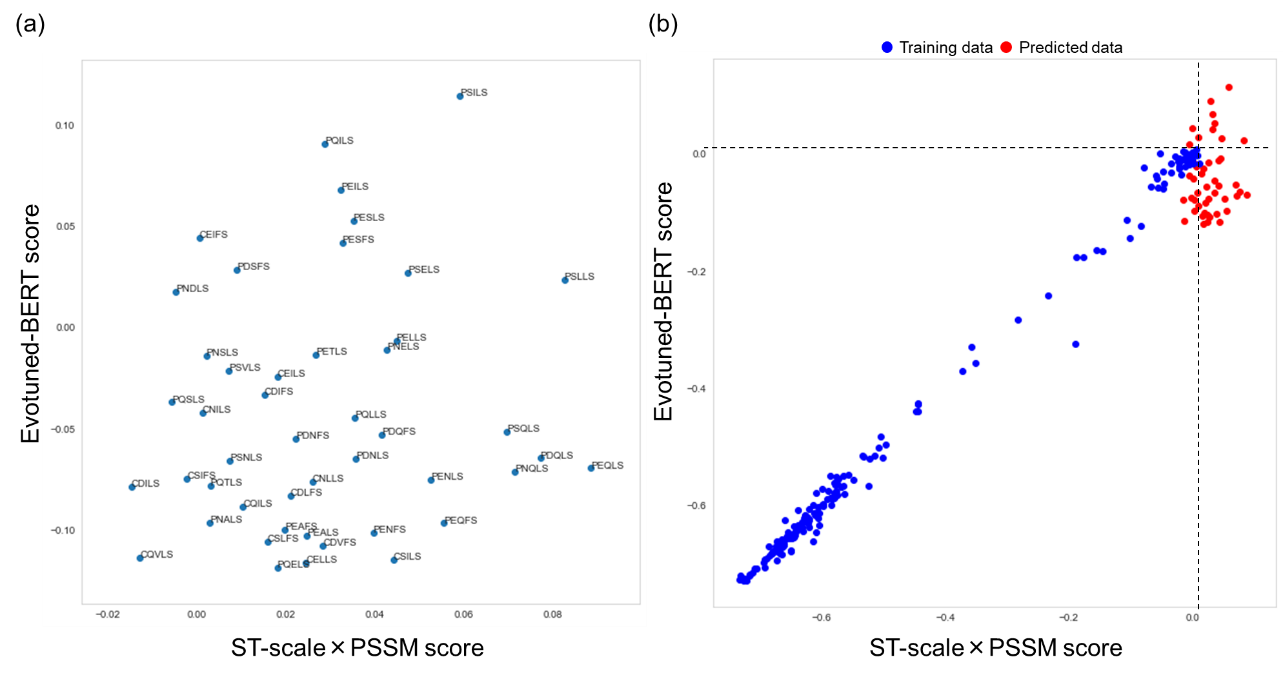

**Figure S6.** Consensus analysis of the second-round top 100 variants predicted by ST-scale×PSSM and Evotuned-BERT models. Predicted scores by the two models are shown. (a) 46 XylM variants commonly predicted by both models to be in the top 100. (b) Comparison of XylM variants of the initial/first-round libraries (training data) and the second-round top 100 prediction. The highest-activity variant in the training data is shown with dashed lines.

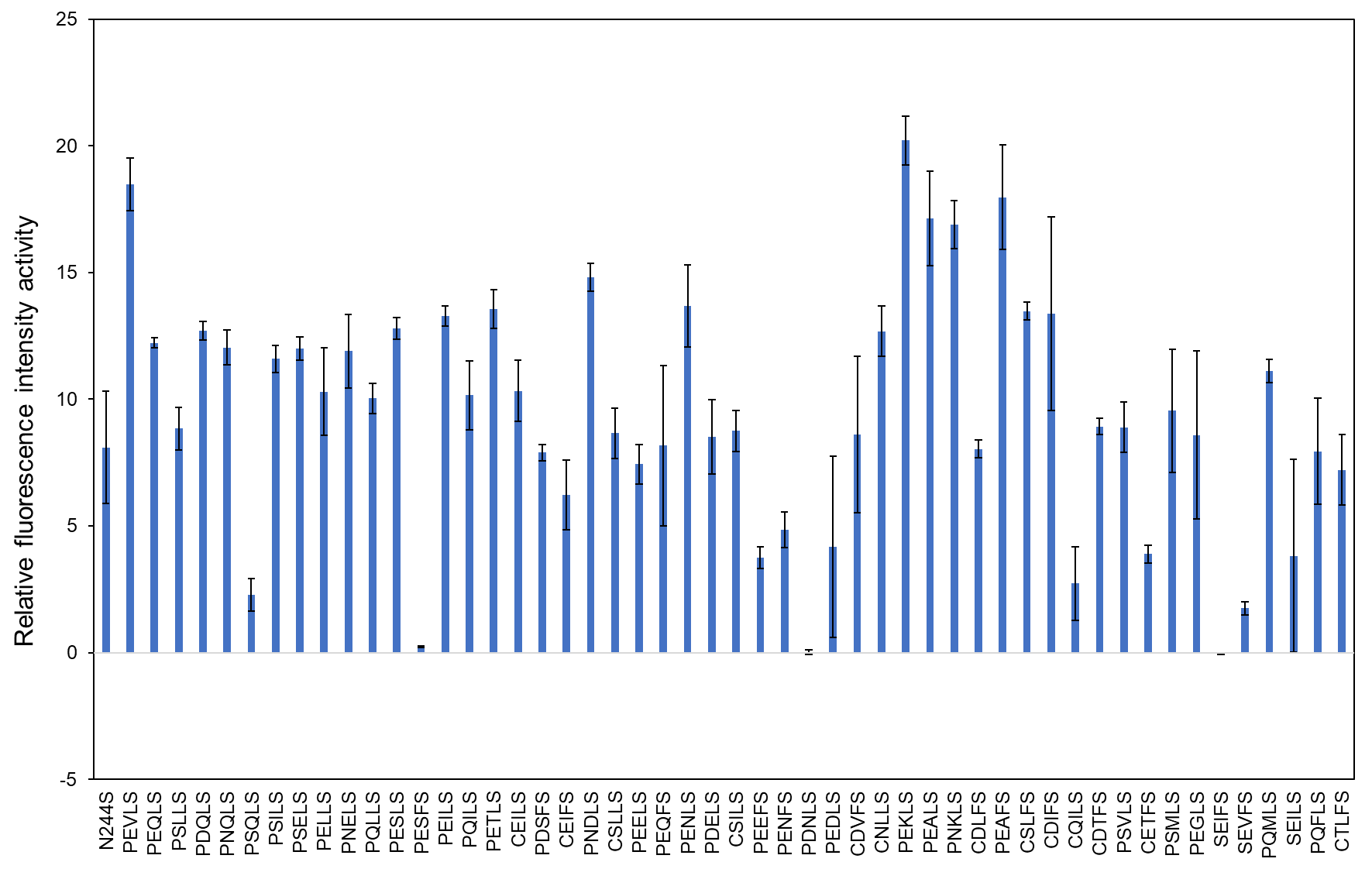

**Figure S7.** The relative fluorescence intensity activities of XylM variants of the second-round library. The error bars indicate standard error (n = 3).

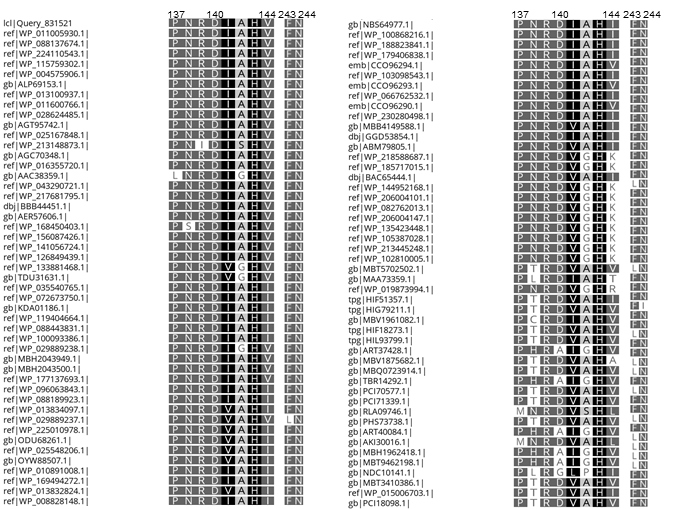

**Figure S8.** Multiple sequence alignment of 96 XylM homologs at P137, D140, V144, F243, and N244. The corresponding amino acids at these residues are shown. These sequences were obtained by NCBI BLAST search using the wild-type XylM sequence as a query. Among the top 100 sequences, 96 sequences covering P137-N244 were selected for the multiple alignment.

**Data S1.** Ranking list by ML prediction using the initial XylM variants library as training data. This list was used for designing the first-round XylM variants library. The top 100,000 variants are shown. The data are available as a separate Excel file.

**Data S2.** Ranking list by ML prediction using the first-round XylM variants library as additional training data. This list was used for designing the second-round XylM variants library. The top 100,000 variants are shown. The data are provided as a separate Excel file.

Table S1 (Underline indicates the restriction sites. Bold letters indicate the mutation sites.)

| No | Primer | Sequence (5’ to 3’) | Use |
| --- | --- | --- | --- |
| 1 | XylMAC-PstI | CCGCTGCAGATGCGGGAAACAAAAGAGCA | Amplification of *xylMABC* |
| 2 | XylMAC-KpnI | GCCGGTACCTCAACCAATCCGGAGTACCG | Amplification of *xylMABC* |
| 3 | XylMA-F | GATGGGTTTATGAATGAGTT | Correction of unintentional mutation introduced by PCR error |
| 4 | XylMA-R | TCAAATGCTAGCCACCCGAC | Correction of unintentional mutation introduced by PCR error |
| 5 | placF-NcoI | GCCCCATGGGCGCAACGCAATTAATGTGA | Amplification of *lac* promoter |
| 6 | placR-PstI | GCCCTGCAGTGTGTTTCCTGTGTGAAATTG | Amplification of *lac* promoter |
| 7 | cat-F | TGATTGAAAAAGGAAGAGTATGGAGAAAAAAATCACTGGA | Amplification of *cat* |
| 8 | cat-R | GTAAACTTGGTCTGACAGTTACGCCCCGCCCTGC | Amplification of *cat* |
| 9 | pETlacXylMAC-F | ACTCTTCCTTTTTCAATCAT | Amplification of pETlacXylMABC-vector |
| 10 | pETlacXylMAC-R | CTGTCAGACCAAGTTTACTC | Amplification of pETlacXylMABC-vector |
| 11 | XylCremove-F | TTCACACAGGAAACACATATGGACACGCTTCGTTATTA | Removal of *xylC* |
| 12 | XylCremove-R | ATGTGTTTCCTGTGTGAAAT | Removal of *xylC* |
| 13 | XylM-seq-F | GCCAATTAACTGTGAGCCGC | Sequence analysis of *xylM* |
| 14 | XylM-seq-R | CCTTGATGCAGAGCGCTTTC | Equence analysis of *xylM* |
| 15 | XylM-D140A-F | GGTGATCCGAACCGA**GCC**ATTGCCCATGTCAACACG | Site-directed mutagenesis of *xylM* to introduce D140A |
| 16 | XylM-D140A-R | TCGGTTCGGATCACCATAAA | Site-directed mutagenesis of *xylM* to introduce D140A |
| 17 | XylM-H143A-F | CGAGACATTGCCCAT**GCC**AACACGCATCACCTTTAC | Site-directed mutagenesis of *xylM* to introduce H143A |
| 18 | XylM-H143A-R | ATGGGCAATGTCTCGGTTCG | Site-directed mutagenesis of *xylM* to introduce H143A |
| 19 | XylM-V144A-F | CGAGACATTGCCCAT**GCC**AACACGCATCACCTTTAC | Site-directed mutagenesis of *xylM* to introduce V144A |
| 20 | XylM-H144A-R | ATGGGCAATGTCTCGGTTCG | Site-directed mutagenesis of *xylM* to introduce V144A |
| 21 | XylM-D156A-F | TTAGATACGCCTCTC**GCC**AGCGATACTCCGTACCGT | Site-directed mutagenesis of *xylM* to introduce D156A |
| 22 | XylM-D156A-R | GAGAGGCGTATCTAAGTAAA | Site-directed mutagenesis of *xylM* to introduce D156A |
| 23 | XylM-D158A-F | ACGCCTCTCGATAGC**GCC**ACTCCGTACCGTGGTCAG | Site-directed mutagenesis of *xylM* to introduce D158A |
| 24 | XylM-D158A-R | GCTATCGAGAGGCGTATCTA | Site-directed mutagenesis of *xylM* to introduce D158A |
| 25 | XylM-Y167A-F | CGTGGTCAGACAATT**GCC**AGTTTCGTGATCAGTGC | Site-directed mutagenesis of *xylM* to introduce Y167A |
| 26 | XylM-Y167A-R | AATTGTCTGACCACGGTACG | Site-directed mutagenesis of xylM to introduce Y167A |
| 27 | XylM-V240A-F | ATTGCGAAAGGGATA**GCC**GAGGGTTTTAATTACTTT | Site-directed mutagenesis of xylM to introduce V240A |
| 28 | XylM-V240A-R | ATCCCTTTCGCAATAATCAT | Site-directed mutagenesis of *xylM* to introduce V240A |
| 29 | XylM-F243A-F | GGGATAGTCGAGGGT**GCC**AATTACTTTCAGCACTAT | Site-directed mutagenesis of *xylM* to introduce F243A |
| 30 | XylM-F243A-R | ACCCTCGACTATCCCTTTCG | Site-directed mutagenesis of *xylM* to introduce F243A |
| 31 | XylM-N244A-F | ATAGTCGAGGGTTTT**GCC**TACTTTCAGCACTATGGT | Site-directed mutagenesis of *xylM* to introduce N244A |
| 32 | XylM-N244A-R | AAAACCCTCGACTATCCCTT | Site-directed mutagenesis of *xylM* to introduce N244A |
| 33 | XylM-H248A-F | TTTAATTACTTTCAG**GCC**TATGGTTTAGTACGCGAT | Site-directed mutagenesis of *xylM* to introduce H248A |
| 34 | XylM-H248A-R | CTGAAAGTAATTAAAACCCT | Site-directed mutagenesis of *xylM* to introduce H248A |
| 35 | XylM-I279A-F | CCGCTGGGTTGCGAA**GCC**ACTAACCATATCAATCAT | Site-directed mutagenesis of *xylM* to introduce I279A |
| 36 | XylM-I279A-R | TTCGCAACCCAGCGGGCGCA | Site-directed mutagenesis of *xylM* to introduce I279A |
| 37 | XylM-H282A-F | ACTAACGCCATCAATCATCATATTGAC | Site-directed mutagenesis of *xylM* to introduce H282A |
| 38 | XylM-H282A-R | ATTGATGGCGTTAGTAATTTCGCAACC | Site-directed mutagenesis of *xylM* to introduce H282A |
| 39 | XylM-H143A-F | AACCGAGACATTGCC**GCC**GTCAACACGCATCACCTT | Site-directed mutagenesis of *xylM* to introduce H143A |
| 40 | XylM-H143A-R | GGCAATGTCTCGGTTCGGAT | Site-directed mutagenesis of *xylM* to introduce H143A |
| 41 | XylM-D136A-F | GCTATGTTTTATGGT**GCC**CCGAACCGAGACATTGCC | Site-directed mutagenesis of *xylM* to introduce D136A |
| 42 | XylM-D136A-R | ACCATAAAACATAGCCAATA | Site-directed mutagenesis of *xylM* to introduce D136A |
| 43 | XylM-P137A-F | ATGTTTTATGGTGAT**GCC**AACCGAGACATTGCCCAT | Site-directed mutagenesis of xylM to introduce P137A |
| 44 | XylM-P137A-R | ATCACCATAAAACATAGCCA | Site-directed mutagenesis of *xylM* to introduce P137A |
| 45 | XylM-L149A-F | GTCAACACGCATCAC**GCC**TACTTAGATACGCCTCTC | Site-directed mutagenesis of *xylM* to introduce L149A |
| 46 | XylM-L149A-R | GTGATGCGTGTTGACATGGG | Site-directed mutagenesis of *xylM* to introduce L149A |
| 47 | XylM-T108A-F | CTTAGTGGTGTGCCA**GCC**CTTCCGGTTTCGCATGAG | Site-directed mutagenesis of *xylM* to introduce T108A |
| 48 | XylM-T108A-R | TGGCACACCACTAAGCCAAG | Site-directed mutagenesis of *xylM* to introduce T108A |
| 49 | XylM-F243N244NNT | GGGATAGTCGAGGGT**NNTNNT**TACTTTCAGCACTATGGT | Site-saturation mutagenesis of *xylM* at F243 and N244. |
| 50 | XylM-N244S | ATAGTCGAGGGTTTT**AGT**TACTTTCAGCACTATGGT | Site-directed mutagenesis of *xylM* to introduce N244S |
| 51 | XylM-F243NTT-N244S | GGGATAGTCGAGGGT**NTTAGT**TACTTTCAGCACTATGGT | Site-saturation mutagenesis of *xylM*-N244S at F243. |
| 52 | XylM-N244NNT | ATAGTCGAGGGTTTT**NNT**TACTTTCAGCACTATGGT | Site-saturation mutagenesis of *xylM* at N244 |
| 53 | XylM-140NNT | GGTGATCCGAACCGA**NNT**ATTGCCCATGTCAACACG | Site-saturation mutagenesis of *xylM* at D140 |
| 54 | XylM-144NNT | CGAGACATTGCCCAT**NNT**AACACGCATCACCTTTAC | Site-saturation mutagenesis of *xylM* at V144 |
| 55 | XylM-243NNT | GGGATAGTCGAGGGT**NNT**AATTACTTTCAGCACTAT | Site-saturation mutagenesis of *xylM* at F243 |
| 56 | XylM-137NNT | ATGTTTTATGGTGAT**NNT**AACCGAGACATTGCCCAT | Site-saturation mutagenesis of *xylM* at P137 |
| 57 | XylM-137MHS | ATGTTTTATGGTGAT**MHS**AACCGAGACATTGCCCAT | Codon-randomization mutagenesis of *xylM* at P137 |
| 58 | XylM-140-MHS | GGTGATCCGAACCGA**MHS**ATTGCCCATGTCAACACG | Codon-randomization mutagenesis of xylM at D140 |
| 59 | XylM-144MHS | CGAGACATTGCCCAT**MHS**AACACGCATCACCTTTAC | Codon-randomization mutagenesis of *xylM* at V144 |
| 60 | XylM-243MHS | GGGATAGTCGAGGGT**MHS**AATTACTTTCAGCACTAT | Codon-randomization mutagenesis of *xylM* at F243 |
| 61 | XylM-244MHS | ATAGTCGAGGGTTTT**MHS**TACTTTCAGCACTATGGT | Codon-randomization mutagenesis of *xylM* at N244 |
| 62 | XylM-137DGG | ATGTTTTATGGTGAT**DGG**AACCGAGACATTGCCCAT | Codon-randomization mutagenesis of *xylM* at P137 |
| 63 | XylM-140-DGG | GGTGATCCGAACCGA**DGG**ATTGCCCATGTCAACACG | Codon-randomization mutagenesis of *xylM* at D140 |
| 64 | XylM-144-DGG | CGAGACATTGCCCAT**DGG**AACACGCATCACCTTTAC | Codon-randomization mutagenesis of *xylM* at V144 |
| 65 | XylM-243-DGG | GGGATAGTCGAGGGT**DGG**AATTACTTTCAGCACTAT | Codon-randomization mutagenesis of *xylM* at F243 |
| 66 | XylM-244-DGG | ATAGTCGAGGGTTTT**DGG**TACTTTCAGCACTATGGT | Codon-randomization mutagenesis of *xylM* at N244 |
| 67 | XylM-137WWC | ATGTTTTATGGTGAT**WWC**AACCGAGACATTGCCCAT | Codon-randomization mutagenesis of *xylM* at P137 |
| 68 | XylM-137E | ATGTTTTATGGTGAT**GAG**AACCGAGACATTGCCCAT | Site-directed mutagenesis of *xylM* to introduce P137E |
| 69 | XylM-140TWC | GGTGATCCGAACCGA**TWC**ATTGCCCATGTCAACACG | Codon-randomization mutagenesis of *xylM* at D140 |
| 70 | XylM-140AWS | GGTGATCCGAACCGA**AWS**ATTGCCCATGTCAACACG | Codon-randomization mutagenesis of *xylM* at D140 |
| 71 | XylM-140E | GGTGATCCGAACCGA**GAG**ATTGCCCATGTCAACACG | Site-directed mutagenesis of *xylM* to introduce D140E |
| 72 | XylM-144ABC | CGAGACATTGCCCAT**ABC**AACACGCATCACCTTTAC | Codon-randomization mutagenesis of *xylM* at V144 |
| 73 | XylM-144GAG | CGAGACATTGCCCAT**GAG**AACACGCATCACCTTTAC | Codon-randomization mutagenesis of *xylM* at V144 |
| 74 | XylM-243TRC | GGGATAGTCGAGGGT**TRC**AATTACTTTCAGCACTAT | Codon-randomization mutagenesis of *xylM* at F243 |
| 75 | XylM-243AVS | GGGATAGTCGAGGGT**AVS**AATTACTTTCAGCACTAT | Codon-randomization mutagenesis of *xylM* at F243 |
| 76 | XylM-243E | GGGATAGTCGAGGGT**GAG**AATTACTTTCAGCACTAT | Site-directed mutagenesis of *xylM* to introduce F243E |
| 77 | XylM-244WTT | ATAGTCGAGGGTTTT**WTT**TACTTTCAGCACTATGGT | Codon-randomization mutagenesis of *xylM* at N244 |
| 78 | XylM-244E | ATAGTCGAGGGTTTT**GAG**TACTTTCAGCACTATGGT | Site-directed mutagenesis of *xylM* to introduce N244E |
| 79 | XylM-137F | ATGTTTTATGGTGAT**TTC**AACCGAGACATTGCCCAT | Site-directed mutagenesis of *xylM* to introduce P137F |
| 80 | XylM-137M | ATGTTTTATGGTGAT**ATG**AACCGAGACATTGCCCAT | Site-directed mutagenesis of *xylM* to introduce P137M |
| 81 | XylM-137W | ATGTTTTATGGTGAT**TGG**AACCGAGACATTGCCCAT | Site-directed mutagenesis of *xylM* to introduce P137W |
| 82 | XylM-140F | GGTGATCCGAACCGA**TTC**ATTGCCCATGTCAACACG | Site-directed mutagenesis of *xylM* to introduce D140F |
| 83 | XylM-140M | GGTGATCCGAACCGA**ATG**ATTGCCCATGTCAACACG | Site-directed mutagenesis of *xylM* to introduce D140M |
| 84 | XylM-144I | CGAGACATTGCCCAT**ATT**AACACGCATCACCTTTAC | Site-directed mutagenesis of *xylM* to introduce V144I |
| 85 | XylM-144T | CGAGACATTGCCCAT**ACC**AACACGCATCACCTTTAC | Site-directed mutagenesis of *xylM* to introduce V144T |
| 86 | XylM-243K | GGGATAGTCGAGGGT**AAA**AATTACTTTCAGCACTAT | Site-directed mutagenesis of *xylM* to introduce F243K |
| 87 | XylM-243N | GGGATAGTCGAGGGT**AAC**AATTACTTTCAGCACTAT | Site-directed mutagenesis of *xylM* to introduce F243N |
| 88 | XylM-243T | GGGATAGTCGAGGGT**ACT**AATTACTTTCAGCACTAT | Site-directed mutagenesis of xylM to introduce F243T |
| 89 | XylM-243Y | GGGATAGTCGAGGGT**TAC**AATTACTTTCAGCACTAT | Site-directed mutagenesis of *xylM* to introduce F243Y |
| 90 | XylM-244F | ATAGTCGAGGGTTTT**TTC**TACTTTCAGCACTATGGT | Site-directed mutagenesis of *xylM* to introduce N244F |
| 91 | XtlM-244P | ATAGTCGAGGGTTTT**CCT**TACTTTCAGCACTATGGT | Site-directed mutagenesis of xylM to introduce N244P |
| 92 | XylM-149R | GTCAACACGCATCAC**CGA**TACTTAGATACGCCTCTC | Site-directed mutagenesis of *xylM* to introduce L149R |
| 93 | XylM-137-244seq | GGTGTGCCAACTCTTCCGGT | Sequence analysis of *xylM* |
| 94 | XylM-137140144-NNT | ATGTTTTATGGTGAT**NNT**AACCGA**NNT**ATTGCCCAT**NNT**AACACGCATCACCTTTAC | Site-saturation mutagenesis of *xylM* at P137, D140, and V144. |
| 95 | XylM-243L244S | GGGATAGTCGAGGGT**TTAAGT**TACTTTCAGCACTATGGT | Site-directed mutagenesis of *xylM* to introduce F243L-N244S |
| 96 | XylM243Y244S | GGGATAGTCGAGGGT**TACAGT**TACTTTCAGCACTATGGT | Site-directed mutagenesis of *xylM* to introduce F243Y- N244S |
| 97 | XylM243M244S | GGGATAGTCGAGGGT**ATGAGT**TACTTTCAGCACTATGGT | Site-directed mutagenesis of *xylM* to introduce F243M- N244S |
| 98 | XylM-144-F | AACACGCATCACCTTTACTT | Amplification of pETlac*XylM*ABC-vector |
| 99 | XylM-PNLFS | TTTATGGTGATCCGAACCGA**AAC**ATTGCCCAT**TTA**AACACGCATCACCTTTACTT | Site-directed mutagenesis of *xylM* at P137, D140, V144, F243, and N244 |
| 100 | XylM-PNIFS | TTTATGGTGATCCGAACCGA**AAC**ATTGCCCAT**ATT**AACACGCATCACCTTTACTT | Site-directed mutagenesis of *xylM* at P137, D140, V144, F243, and N244 |
| 101 | XylM-PELFS | TTTATGGTGATCCGAACCGA**GAG**ATTGCCCAT**TTA**AACACGCATCACCTTTACTT | Site-directed mutagenesis of *xylM* at P137, D140, V144, F243, and N244 |
| 102 | XylM-PEIFS | TTTATGGTGATCCGAACCGA**GAG**ATTGCCCAT**ATT**AACACGCATCACCTTTACTT | Site-directed mutagenesis of *xylM* at P137, D140, V144, F243, and N244 |
| 103 | XylM-PQIFS | TTTATGGTGATCCGAACCGA**CAA**ATTGCCCAT**ATT**AACACGCATCACCTTTACTT | Site-directed mutagenesis of *xylM* at P137, D140, V144, F243, and N244 |
| 104 | XylM-PSLFS | TTTATGGTGATCCGAACCGA**AGT**ATTGCCCAT**CTT**AACACGCATCACCTTTACTT | Site-directed mutagenesis of *xylM* at P137, D140, V144, F243, and N244 |
| 105 | XylM-PSEFS | TTTATGGTGATCCGAACCGA**AGT**ATTGCCCAT**GAG**AACACGCATCACCTTTACTT | Site-directed mutagenesis of *xylM* at P137, D140, V144, F243, and N244 |
| 106 | XylM-PETFS | TTTATGGTGATCCGAACCGA**GAG**ATTGCCCAT**ACG**AACACGCATCACCTTTACTT | Site-directed mutagenesis of *xylM* at P137, D140, V144, F243, and N244 |
| 107 | XylM-PNDFS | TTTATGGTGATCCGAACCGA**AAC**ATTGCCCAT**GAT**AACACGCATCACCTTTACTT | Site-directed mutagenesis of *xylM* at P137, D140, V144, F243, and N244 |
| 108 | XylM-PTDFS | TTTATGGTGATCCGAACCGA**ACG**ATTGCCCAT**GAT**AACACGCATCACCTTTACTT | Site-directed mutagenesis of *xylM* at P137, D140, V144, F243, and N244 |
| 109 | XylM-PSIFS | TTTATGGTGATCCGAACCGA**AGT**ATTGCCCAT**ATT**AACACGCATCACCTTTACTT | Site-directed mutagenesis of *xylM* at P137, D140, V144, F243, and N244 |
| 110 | XylM-PEGFS | TTTATGGTGATCCGAACCGA**GAA**ATTGCCCAT**GGT**AACACGCATCACCTTTACTT | Site-directed mutagenesis of *xylM* at P137, D140, V144, F243, and N244 |
| 111 | XylM-CEVFS | TGGCTATGTTTTATGGTGAT**TGC**AACCGA**GAA**ATTGCCCATGTCAACACGCA | Site-directed mutagenesis of *xylM* at P137, D140, V144, F243, and N244 |
| 112 | XylM-CNVFS | TGGCTATGTTTTATGGTGAT**TGC**AACCGA**AAC**ATTGCCCATGTCAACACGCA | Site-directed mutagenesis of *xylM* at P137, D140, V144, F243, and N244 |
| 113 | XylM-CNLFS | TGGCTATGTTTTATGGTGAT**TGC**AACCGA**AAC**ATTGCCCAT**CTA**AACACGCATCACCTTTACTT | Site-directed mutagenesis of *xylM* at P137, D140, V144, F243, and N244 |
| 114 | XylM-CELFS | TGGCTATGTTTTATGGTGAT**TGC**AACCGA**GAG**ATTGCCCAT**CTA**AACACGCATCACCTTTACTT | Site-directed mutagenesis of *xylM* at P137, D140, V144, F243, and N244 |
| 115 | XylM-CQVFS | TGGCTATGTTTTATGGTGAT**TGC**AACCGA**CAG**ATTGCCCATGTCAACACGCA | Site-directed mutagenesis of *xylM* at P137, D140, V144, F243, and N244 |
| 116 | XylM-PSDLS | TTTATGGTGATCCGAACCGA**AGT**ATTGCCCAT**GAT**AACACGCATCACCTTTACTT | Site-directed mutagenesis of *xylM* at P137, D140, V144, F243, and N244 |
| 117 | XylM-EPVFS | TGGCTATGTTTTATGGTGAT**GAA**AACCGA**CCG**ATTGCCCATGTCAACACGCA | Site-directed mutagenesis of *xylM* at P137, D140, V144, F243, and N244 |
| 118 | XylM-DPVFS | TGGCTATGTTTTATGGTGAT**GAT**AACCGA**CCG**ATTGCCCATGTCAACACGCA | Site-directed mutagenesis of *xylM* at P137, D140, V144, F243, and N244 |
| 119 | XylM-DPLFS | TGGCTATGTTTTATGGTGAT**GAT**AACCGA**CCG**ATTGCCCAT**CTA**AACACGCATCACCTTTACTT | Site-directed mutagenesis of *xylM* at P137, D140, V144, F243, and N244 |
| 120 | XylM-EPLFS | TGGCTATGTTTTATGGTGAT**GAG**AACCGA**CCG**ATTGCCCAT**CTA**AACACGCATCACCTTTACTT | Site-directed mutagenesis of *xylM* at P137, D140, V144, F243, and N244 |
| 121 | XylM-SPEFS | TGGCTATGTTTTATGGTGAT**AGT**AACCGA**CCG**ATTGCCCAT**GAG**AACACGCATCACCTTTACTT | Site-directed mutagenesis of *xylM* at P137, D140, V144, F243, and N244 |
| 122 | XylM-SPDFS | TGGCTATGTTTTATGGTGAT**AGT**AACCGA**CCG**ATTGCCCAT**GAT**AACACGCATCACCTTTACTT | Site-directed mutagenesis of *xylM* at P137, D140, V144, F243, and N244 |
| 123 | XylM-SDIFS | TGGCTATGTTTTATGGTGAT**AGT**AACCGA**GAT**ATTGCCCAT**ATT**AACACGCATCACCTTTACTT | Site-directed mutagenesis of *xylM* at P137, D140, V144, F243, and N244 |
| 124 | XylM-SDFFS | TGGCTATGTTTTATGGTGAT**AGT**AACCGA**GAT**ATTGCCCAT**TTT**AACACGCATCACCTTTACTT | Site-directed mutagenesis of *xylM* at P137, D140, V144, F243, and N244 |
| 125 | XylM-APDFS | TGGCTATGTTTTATGGTGAT**GCT**AACCGA**CCC**ATTGCCCAT**GAT**AACACGCATCACCTTTACTT | Site-directed mutagenesis of *xylM* at P137, D140, V144, F243, and N244 |
| 126 | XylM-140-F | ATTGCCCATGTCAACACGCA | Amplification of pETlac*XylM*ABC-vector |
| 127 | XylM-140R | GGTGATCCGAACCGA**CGA**ATTGCCCATGTCAACACG | Amplification of pETlac*XylM*ABC-vector |
| 128 | XylM-PEQLS | TTTATGGTGATCCGAACCGA**GAA**ATTGCCCAT**CAG**AACACGCATCACCTTTACTT | Site-directed mutagenesis of *xylM* at P137, D140, V144, F243, and N244 |
| 129 | XylM-PNQLS | TTTATGGTGATCCGAACCGA**AAC**ATTGCCCAT**CAG**AACACGCATCACCTTTACTT | Site-directed mutagenesis of *xylM* at P137, D140, V144, F243, and N244 |
| 130 | XylM-PSQLS | TTTATGGTGATCCGAACCGA**AGC**ATTGCCCAT**CAG**AACACGCATCACCTTTACTT | Site-directed mutagenesis of *xylM* at P137, D140, V144, F243, and N244 |
| 131 | XylM-PNELS | TTTATGGTGATCCGAACCGA**AAC**ATTGCCCAT**GAA**AACACGCATCACCTTTACTT | Site-directed mutagenesis of *xylM* at P137, D140, V144, F243, and N244 |
| 132 | XylM-PQLLS | TTTTATGGTGATCCGAACCGA**CAG**ATTGCCCAT**TTA**AACACGCATCACCTTTACT | Site-directed mutagenesis of *xylM* at P137, D140, V144, F243, and N244 |
| 133 | XylM-PESLS | TTTATGGTGATCCGAACCGA**GAA**ATTGCCCAT**AGC**AACACGCATCACCTTTACTT | Site-directed mutagenesis of *xylM* at P137, D140, V144, F243, and N244 |
| 134 | XylM-CEILS | TGGCTATGTTTTATGGTGAT**TGC**AACCGA**GAA**ATTGCCCAT**ATT**AACACGCATCACCTTTACTT | Site-directed mutagenesis of *xylM* at P137, D140, V144, F243, and N244 |
| 135 | XylM-CSLLS | TGGCTATGTTTTATGGTGAT**TGC**AACCGA**AGC**ATTGCCCAT**TTA**AACACGCATCACCTTTACTT | Site-directed mutagenesis of *xylM* at P137, D140, V144, F243, and N244 |
| 136 | XylM-PEELS | TTTATGGTGATCCGAACCGA**GAA**ATTGCCCAT**GAA**AACACGCATCACCTTTACTT | Site-directed mutagenesis of *xylM* at P137, D140, V144, F243, and N244 |
| 137 | XylM-PENLS | TTTATGGTGATCCGAACCGA**GAA**ATTGCCCAT**AAC**AACACGCATCACCTTTACTT | Site-directed mutagenesis of *xylM* at P137, D140, V144, F243, and N244 |
| 138 | XylM-CSILS | TGGCTATGTTTTATGGTGAT**TGC**AACCGA**AGC**ATTGCCCAT**ATT**AACACGCATCACCTTTACTT | Site-directed mutagenesis of *xylM* at P137, D140, V144, F243, and N244 |
| 139 | XylM-PEDLS | TTTATGGTGATCCGAACCGA**GAA**ATTGCCCAT**GAT**AACACGCATCACCTTTACTT | Site-directed mutagenesis of *xylM* at P137, D140, V144, F243, and N244 |
| 140 | XylM-PEKLS | TTTATGGTGATCCGAACCGA**GAA**ATTGCCCAT**AAA**AACACGCATCACCTTTACTT | Site-directed mutagenesis of *xylM* at P137, D140, V144, F243, and N244 |
| 141 | XylM-PEALS | TTTATGGTGATCCGAACCGA**GAA**ATTGCCCAT**GCT**AACACGCATCACCTTTACTT | Site-directed mutagenesis of *xylM* at P137, D140, V144, F243, and N244 |
| 142 | XylM-PNKLS | TTTATGGTGATCCGAACCGA**AAC**ATTGCCCAT**AAA**AACACGCATCACCTTTACTT | Site-directed mutagenesis of *xylM* at P137, D140, V144, F243, and N244 |
| 143 | XylM-CQILS | TGGCTATGTTTTATGGTGAT**TGC**AACCGA**CAG**ATTGCCCAT**ATT**AACACGCATCACCTTTACTT | Site-directed mutagenesis of *xylM* at P137, D140, V144, F243, and N244 |
| 144 | XylM-PSMLS | TTTATGGTGATCCGAACCGA**AGC**ATTGCCCAT**ATG**AACACGCATCACCTTTACTT | Site-directed mutagenesis of *xylM* at P137, D140, V144, F243, and N244 |
| 145 | XylM-SEIFS | TGGCTATGTTTTATGGTGAT**AGC**AACCGA**GAA**ATTGCCCAT**ATT**AACACGCATCACCTTTACTT | Site-directed mutagenesis of *xylM* at P137, D140, V144, F243, and N244 |
| 146 | XylM-PQMLS | TTTATGGTGATCCGAACCGA**CAG**ATTGCCCAT**ATG**AACACGCATCACCTTTACTT | Site-directed mutagenesis of *xylM* at P137, D140, V144, F243, and N244 |
| 147 | XylM-PQFLS | TTTATGGTGATCCGAACCGA**CAG**ATTGCCCAT**TTT**AACACGCATCACCTTTACTT | Site-directed mutagenesis of *xylM* at P137, D140, V144, F243, and N244 |
| 148 | XylM-CDLFS | TGGCTATGTTTTATGGTGAT**TGC**AACCGA**GAC**ATTGCCCATTTAAACACGCATCACCTTTACTT | Site-directed mutagenesis of *xylM* at P137, D140, V144, F243, and N244 |
| 149 | XylM-CDIFS | TGGCTATGTTTTATGGTGAT**TGC**AACCGA**GAC**ATTGCCCAT**ATT**AACACGCATCACCTTTACTT | Site-directed mutagenesis of *xylM* at P137, D140, V144, F243, and N244 |
| 150 | XylM-CDTFS | TGGCTATGTTTTATGGTGAT**TGC**AACCGA**GAC**ATTGCCCAT**ACG**AACACGCATCACCTTTACTT | Site-directed mutagenesis of *xylM* at P137, D140, V144, F243, and N244 |
| 151 | XylM-CETFS | TGGCTATGTTTTATGGTGAT**TGC**AACCGA**GAA**ATTGCCCAT**ACG**AACACGCATCACCTTTACTT | Site-directed mutagenesis of *xylM* at P137, D140, V144, F243, and N244 |
| 152 | XylM-SEVFS | TGGCTATGTTTTATGGTGAT**AGC**AACCGA**GAA**ATTGCCCAT**GTC**AACACGCA | Site-directed mutagenesis of *xylM* at P137, D140, V144, F243, and N244 |
| 153 | XylM-CTLFS | TGGCTATGTTTTATGGTGAT**TGC**AACCGA**ACG**ATTGCCCAT**TTA**AACACGCATCACCTTTACTT | Site-directed mutagenesis of *xylM* at P137, D140, V144, F243, and N244 |
